# GIPC3 couples to MYO6 and PDZ domain proteins and shapes the hair cell apical region

**DOI:** 10.1101/2023.02.28.530466

**Authors:** Paroma Chatterjee, Clive P. Morgan, Jocelyn F. Krey, Connor Benson, Jennifer Goldsmith, Michael Bateschell, Anthony J. Ricci, Peter G. Barr-Gillespie

## Abstract

GIPC3 has been implicated in auditory function. Initially localized to the cytoplasm of inner and outer hair cells of the cochlea, GIPC3 increasingly concentrated in cuticular plates and at cell junctions during postnatal development. Early postnatal *Gipc3^KO/KO^* mice had mostly normal mechanotransduction currents, but had no auditory brainstem response at one month of age. Cuticular plates of *Gipc3^KO/KO^* hair cells did not flatten during development as did those of controls; moreover, hair bundles were squeezed along the cochlear axis in mutant hair cells. Junctions between inner hair cells and adjacent inner phalangeal cells were also severely disrupted in *Gipc3^KO/KO^* cochleas. GIPC3 bound directly to MYO6, and the loss of MYO6 led to altered distribution of GIPC3. Immunoaffinity purification of GIPC3 from chicken inner ear extracts identified co-precipitating proteins associated with adherens junctions, intermediate filament networks, and the cuticular plate. Several of immunoprecipitated proteins contained GIPC-family consensus PDZ binding motifs (PBMs), including MYO18A, which binds directly to the PDZ domain of GIPC3. We propose that GIPC3 and MYO6 couple to PBMs of cytoskeletal and cell-junction proteins to shape the cuticular plate.

**Summary statement:** The PDZ-domain protein GIPC3 couples the molecular motors MYO6 and MYO18A to actin cytoskeleton structures in hair cells. GIPC3 is necessary for shaping the hair cell’s cuticular plate and hence the arrangement of the stereocilia in the hair bundle.

## Introduction

The PDZ (PSD95, DLG1, ZO1) protein GIPC3 is located in sensory hair cells and spiral ganglion neurons and is necessary for auditory function in mice (Charizopoulou et al., 2011) and humans (Charizopoulou et al., 2011, Rehman et al., 2011). GIPC3 belongs to the GIPC family, which also includes GIPC1 and GIPC2; GIPC proteins interact with key transmembrane proteins like receptor tyrosine kinases, G-protein-coupled receptors, and integrins, and play roles in their trafficking, signaling, and internalization (Katoh, 2013). GIPC proteins contain a single PDZ domain that is flanked by a N-terminal GIPC-homology 1 (GH1) domain and a C-terminal GH2 domain (Katoh, 2013). In the absence of ligand, GIPC1 resides in an autoinhibited configuration, where sites that interact with other proteins are masked (Shang et al., 2017). The GIPC1 PDZ domain binds to many PDZ-binding motifs (PBMs), which are typically C-terminal tails of interacting proteins (Katoh, 2013); binding to PBMs then releases autoinhibition, allowing GH2 to bind to other proteins (Shang et al., 2017).

MYO6 is a well characterized interacting partner for GIPC1 (Naccache et al., 2006, Reed et al., 2005, Shang et al., 2017). MYO6 is an unconventional myosin motor that is highly expressed in hair cells and is essential for their function (Avraham et al., 1995, Avraham et al., 1997, Hasson et al., 1997); preliminary evidence suggests that MYO6 also interacts with GIPC3 (Shang et al., 2017). MYO6 interacts with GH2 of GIPC1 (Naccache et al., 2006), so when proteins with PBMs bind to the GIPC1 PDZ domain, the GIPC1 GH2 domain is released and can bind to MYO6.

The original *ahl5* allele identified for *Gipc3*, which contains a missense mutation in the PDZ domain, had elevated auditory brainstem response (ABR) thresholds but not profound deafness (Charizopoulou et al., 2011). By contrast, the International Mouse Phenotyping Consortium (IMPC) (Brown et al., 2018) reported that a full knockout of *Gipc3* was profoundly deaf (www.mousephenotype.org), prompting us to examine GIPC3’s location and role in hair-cell function in more detail.

We developed monoclonal antibodies against GIPC3; we used them to reveal that GIPC3 was initially found in the cytoplasm of cochlear inner hair cells (IHCs), then increasingly concentrated at cell-cell junctions (Corwin and Warchol, 1991) and the cuticular plate (Pollock and McDermott, 2015) as development proceeded. Using the IMPC *Gipc3* knockout mouse line, we found that cuticular plates of *Gipc3^KO/KO^* hair cells were significantly disrupted, appearing rounder than those of controls, and cell-cell junctions were significantly disordered; *Gipc3^KO/KO^* mice also had stereocilia organization defects. Later in development, stereocilia begin to fuse together in *Gipc3^KO/KO^* hair cells, eventually producing giant stereocilia. This phenotype resembled the phenotype of *Myo6^KO/KO^* hair cells (Self et al., 1999), suggesting that GIPC3 and MYO6 may be in the same pathway; indeed, GIPC3 and MYO6 interacted directly. Although MYO6 was not mislocalized in *Gipc3^KO/KO^* hair cells, GIPC3 no longer localized to cuticular plates and junctional regions of *Myo6^KO/KO^* hair cells. One of the anti-GIPC3 monoclonal antibodies was used to immunoaffinity purify GIPC3 and its complexes, and the results suggested that GIPC3 associated with actin-rich networks that are associated with cell junctions, including those of MYO6, MYH9, MYH10, and MYO18A. Several co-immunoprecipitated proteins were predicted to have high-affinity PBM interactions with the GIPC family, including APPL2, MYO18A, ACTN1, and ACTN4. We demonstrated that the GH2 domain of GIPC3 interacts with MYO6, and that the PDZ domain of GIPC3 interacts with MYO18A though its C-terminal PDZ-binding domains. We propose that GIPC3 interacts with MYO6 to couple apical junctions to the cuticular plate, which is necessary for proper anchoring of stereocilia.

## Results

### GIPC3 is enriched in hair cells and localizes to apical regions

We detected GIPC3 in hair bundles in shotgun mass spectrometry experiments that characterized proteins of mouse inner-ear samples; GIPC3 was enriched in bundles as compared to the whole epithelium (Krey et al., 2015). In *Pou4f3-Gfp* mice (Krey et al., 2018), which express GFP exclusively in hair cells (Scheffer et al., 2015), GIPC3 was readily detected in GFP-positive cells using data-dependent acquisition (DDA) with FACS-sorted cells; GIPC3 was present both in cochlear (Fig. 1A) and utricle (Fig. 1B) hair cells, but was at low levels in GFP-negative cells. These results showed that GIPC3 was highly enriched in hair cells.

**Figure 1.**
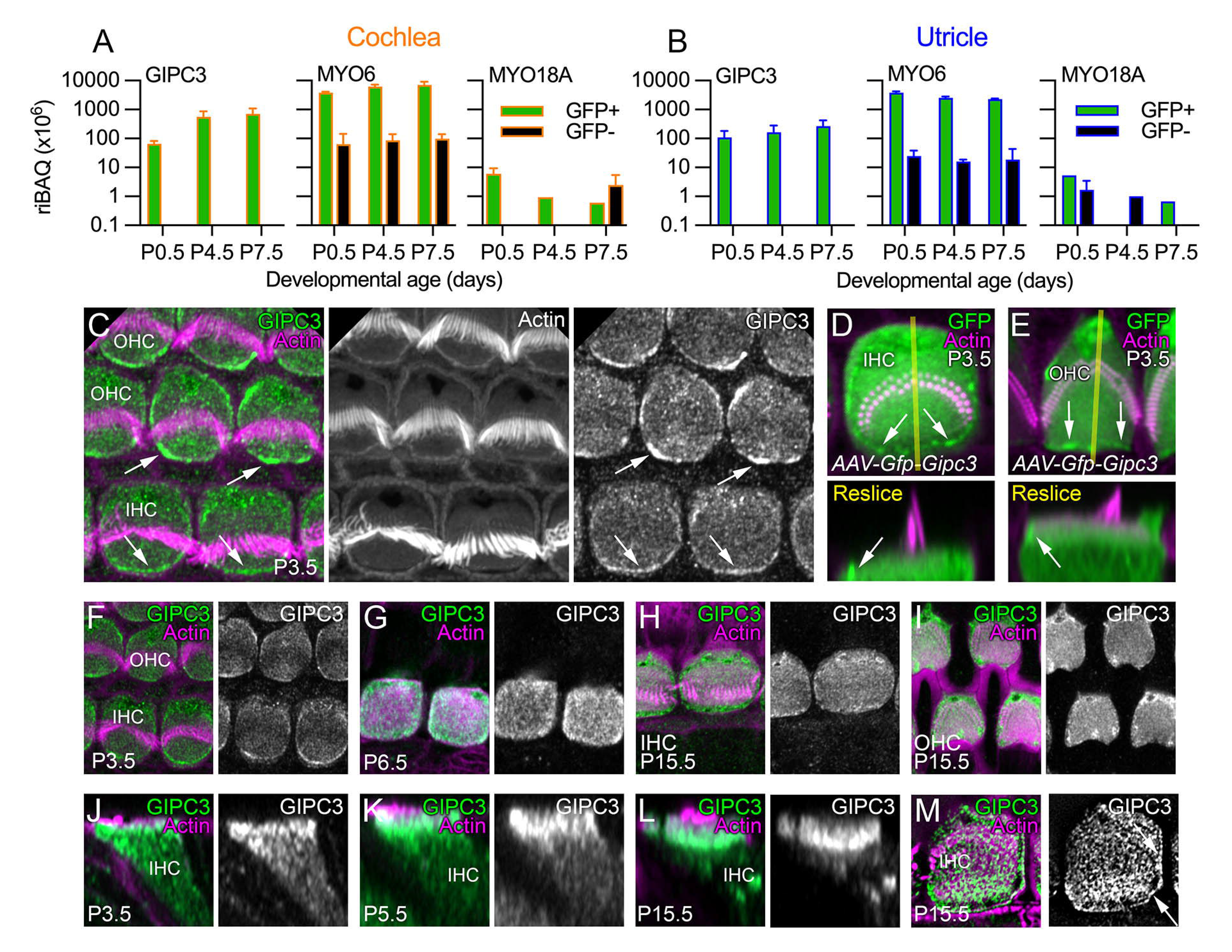
GIPC3 in the cochlea and utricle. ***A***, Mass spectrometry quantitation from cochlea using DDA. For A and B, green corresponds to *Pou4f3*-GFP-positive cells (hair cells), while black corresponds to GFP-negative cells; riBAQ measures relative molar abundance, and mean ± range are plotted (n=2 each). ***B***, Mass spectrometry quantitation from utricle using DDA. ***C***, Localization of GIPC3 in P3.5 cochlear hair cells. OHC, outer hair cell. IHC, inner hair cell. Arrows point to concentration of GIPC3 at the periphery of the cuticular plate, at or adjacent to the plasma membrane in the pericuticular necklace region. ***D-E***, Localization of GFP-GIPC3 introduced into inner (D) and outer (E) hair cells using AAV. Reslice images below are the x-z images from the transects shown in yellow in the upper panels. Arrows point to concentration of GFP-GIPC3 at the pericuticular necklace region. ***F-M***, Developmental progression of GIPC3 labeling in IHCs (F-H, J-M) and OHCs (I). M is a lattice SIM image; arrows indicate GIPC3 located at circumferential actin ring of displayed IHC. AAV experiments were performed twice; immunolocalization experiments were performed >5 times. Panel widths: C, 17.5 µm; D-E, 10 µm; F-I, 15 µm; J-M, 10 µm. Superresolution modality: C-L, Airyscan; M, lattice SIM.

We developed monoclonal antibodies against GIPC3, immunizing mice with a mixture of recombinant mouse and chicken GIPC3 proteins; we chose antibodies designated as 6B4, 3A7, and 10G5 for further characterization of GIPC3. Using these antibodies, which were validated in *Gipc3*-null cochleas (see below), GIPC3 was detected in the cytoplasm of IHCs and OHCs and was modestly enriched in the cuticular plate at early developmental ages; it was particularly concentrated near apical junctions and the circumferential actin belt (arrows in Fig. 1C), in a region called the pericuticular necklace (Hasson et al., 1997). When introduced into hair cells using adeno-associated virus (AAV) transduction via in utero electroporation, GFP-GIPC3 localized to similar locations (Fig. 1D); its concentration at apical junctions was notable (arrows in Fig. 1D-E). GIPC3 detected by any of the three antibodies was found in cuticular plates and at apical junctions throughout development (Figs. 1F-I, S2) but was increasingly concentrated apically by P15.5 (Fig. 1J-L). Lattice structured illumination microscopy (SIM) showed a punctate pattern in the cuticular plate and strong labeling at apical junctions (Fig. 1M, arrows).

### *Gipc3^KO/KO^* have reduced auditory function

We permanently deleted *Gipc3* exons 2 and 3 and the neomycin cassette from *Gipc3^tm1a(KOMP)Wtsi^* mice using the Cre deleter strain (Schwenk et al., 1995); here we refer to the resulting tm1b allele, a global *Gipc3* knockout, as *Gipc3^KO^*. Signal for monoclonal antibodies 6B4 and 3A7 was lost completely in *Gipc3^KO/KO^* cochlea and utricle hair cells (Figs. S1A-F). Most 10G5 signal was lost in *Gipc3^KO/KO^* mice; a very low level of immunoreactivity remained in the junctional region (Fig. S1G-H) and resembled that of GIPC1 immunostaining (Giese et al., 2012). IMPC reported elevated auditory brainstem response (ABR) measurements (www.mousephenotype.org); we replicated those results, showing not only that *Gipc3^KO/KO^* mice were profoundly deaf at 5-6 weeks of age, but that there was a modest threshold elevation in heterozygotes (Fig. 2A).

**Figure 2.**
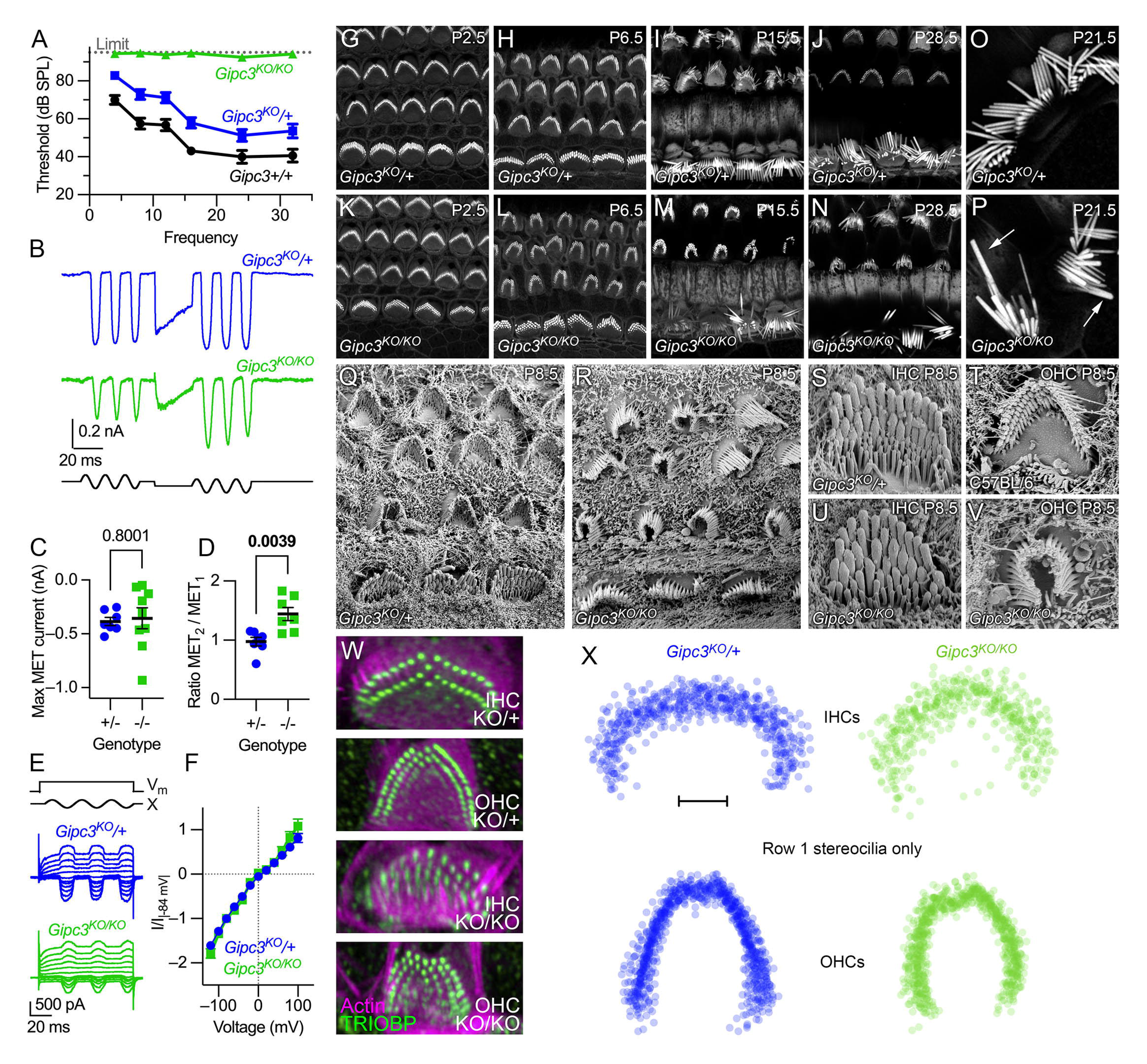
Characterization of *Gipc3* knockout mice. ***A***, ABR thresholds for *Gipc3* genotypes at 5-6 weeks (mean ± s.e.m.). *Gipc3^KO/KO^* completely lacked responses, preventing statistical testing. Unpaired t-test *P*-values for *Gipc3^KO^*/+ comparison to *Gipc3*+/+ were: 4 kHz, 0.0045; 8 kHz, 0.0061; 12 kHz, 0.0207; 16 kHz, 0.0103; 24 kHz, 0.1097; 32 kHz, 0.0751. Sample sizes (n) were (respectively *Gipc3*+/+, *Gipc3^KO^*/+, and *Gipc3^KO/KO^*): 4 kHz, 8 kHz, 16 kHz, 24 kHz, and 32 kHz (8, 28, 18); 12 kHz and 24 kHz (6, 26, 8). ***B-F***, Mechanoelectrical transduction from P10 IHCs; holding potential of -84 mV. ***B***, MET in response to sinusoidal displacement bursts, interceded with a positive adapting step. ***C***, Maximum MET current. Sample sizes were n=7 (*Gipc3^KO^*/+) or 9 (*Gipc3^KO/KO^*). ***D***, Ratio of average responses from second sinusoidal burst divided by responses from first burst (n=7 each). ***E***, MET currents at different voltages; voltage steps ranged from -120 mV to 100 mV in steps of 20 mV. ***F***, MET current-voltage relationships. Current at each voltage was divided by the absolute value of the current at -84 mV (mean ± s.d.). Sample sizes were n=7 (*Gipc3^KO^*/+) or n=4 (*Gipc3^KO/KO^*). ***G-N***, Bundle morphology visualized by phalloidin staining in cochlear hair cells of *Gipc3^KO^*/+ (G-J) and *Gipc3^KO/KO^* (K-M) mice during development. ***O-P***, High magnification view of IHC bundles. Note elongated and thickened *Gipc3^KO/KO^* stereocilia in P (arrows), presumably arising from fusion of several stereocilia. ***Q-R***, Scanning electron microscopy of *Gipc3^KO^*/+ (Q) and *Gipc3^KO/KO^* (R) cochleas. ***S-V***, Magnified views of IHC (S, U) and OHC (T, V) bundles. *Gipc3^KO/KO^* IHC bundles had increased numbers of rows with thick stereocilia (U); *Gipc3^KO/KO^* OHC bundles were often squeezed inwards (V). ***W***, Examples of TRIOBP labeling to detect rootlets of *Gipc3^KO^* heterozygote and knockout IHCs and OHCs. ***X***, Overlays of row 1 stereocilia for *Gipc3^KO^* heterozygote and knockout IHCs and OHCs. Both IHCs and OHC row 1 patterns show inward squeezing. Scale bar in X is 2 µm. Panel widths: G-N, 40 µm; O-P, 20 µm; Q-R, 17.3 µm; S-V, 5 µm; W, 10 µm.

We found that mechanoelectrical transduction (MET) currents at P10 were of similar maximum amplitudes in *Gipc3^KO^*/+ and *Gipc3^KO/KO^* hair cells (Fig. 2B-C). Given the increased number of stereocilia rows, we expected larger currents from *Gipc3^KO/KO^*, but noticed that the fluid jet was less effective at moving their hair bundles. Larger stimulations led to separation of rows and loss of MET currents. By using an offset stimulus, we were able to evoke larger currents from the *Gipc3^KO/KO^* bundles without causing damage. These data were quantified as the ratio of maximum currents after the offset stimulus to that before the stimulus (Fig. 2B, D). The larger currents suggest that the additional rows of stereocilia were functional, albeit harder to stimulate. Application of mechanical stimuli while the cell was subjected to voltage steps allowed us to construct current-voltage relationships for *Gipc3^KO^*/+ and *Gipc3^KO/KO^* MET (Fig. 2E); I-V curves for the two genotypes did not differ (Fig. 2F), suggesting that the MET channel properties were unaffected by the lack of GIPC3.

FM1-43 dye labeling is often used as a proxy for mechanotransduction (Meyers et al., 2003). Dye loading was reduced by about 65% in wild-type littermates using the transduction channel blocker tubocurarine, consistent with most dye entering hair cells via the transduction channels; the reduction was less (∼50%) in heterozygotes, and dye labeling in *Gipc3^KO/KO^* hair cells was only slightly reduced (Figs. S1I-O). These results are consistent with an increased number of transduction channels and an increased open probability at rest.

### Distorted hair bundles in *Gipc3^KO/KO^* inner and outer hair cells

We examined morphology of apical IHCs in *Gipc3^KO/KO^* mice. While hair bundles of P2.5 *Gipc3^KO/KO^* mutant hair cells resembled those of heterozygous controls (Fig. 2G, K), by P6.5 bundles appeared to be squeezed along the cochlear longitudinal axis (Fig. 2H, L). This compression was also apparent by scanning electron microscopy at P8.5 (Fig. 2Q-R). As development progressed, bundles became more disorganized and abnormal (Fig. 2I-J, M-N); after P14.5, fused and elongated stereocilia were often seen (Fig. 2P), resembling those seen in *Myo6* null mutants (Self et al., 1999).

IHC stereocilia were gathered into a wedge shape, rather than a curled block, and diameters of row 3 stereocilia were greater than those in control bundles (Fig. 2S, U). OHC stereocilia shapes were similar in *Gipc3^KO/KO^* and controls, but the wings of the OHC bundles were considerably closer together in the mutants (Fig. 2T, V). TRIOBP labeling (Fig. 2W) also demonstrated that the squeezing distortion seen in bundles was due to altered positions of the stereocilia rootlets; rootlets of row 1 stereocilia in both IHCs and OHCs were reproducibly closer together (Fig. 2X).

### GIPC3 interacts directly with MYO6

Because the paralog GIPC1 directly binds to MYO6, we examined whether GIPC3 also interacts with MYO6. MYO6 was present in cochlear and vestibular hair cells at levels well above those of GIPC3 (Fig. 1A-B). In immunolocalization experiments with IHCs from P6.5 and P15.5 C57BL/6 mice, MYO6 and GIPC3 had overlapping distributions, although MYO6 was relatively more concentrated at apical junctions and the pericuticular region and GIPC3 was more concentrated in the cuticular plate, especially at later ages (Fig. 3A-B).

**Figure 3.**
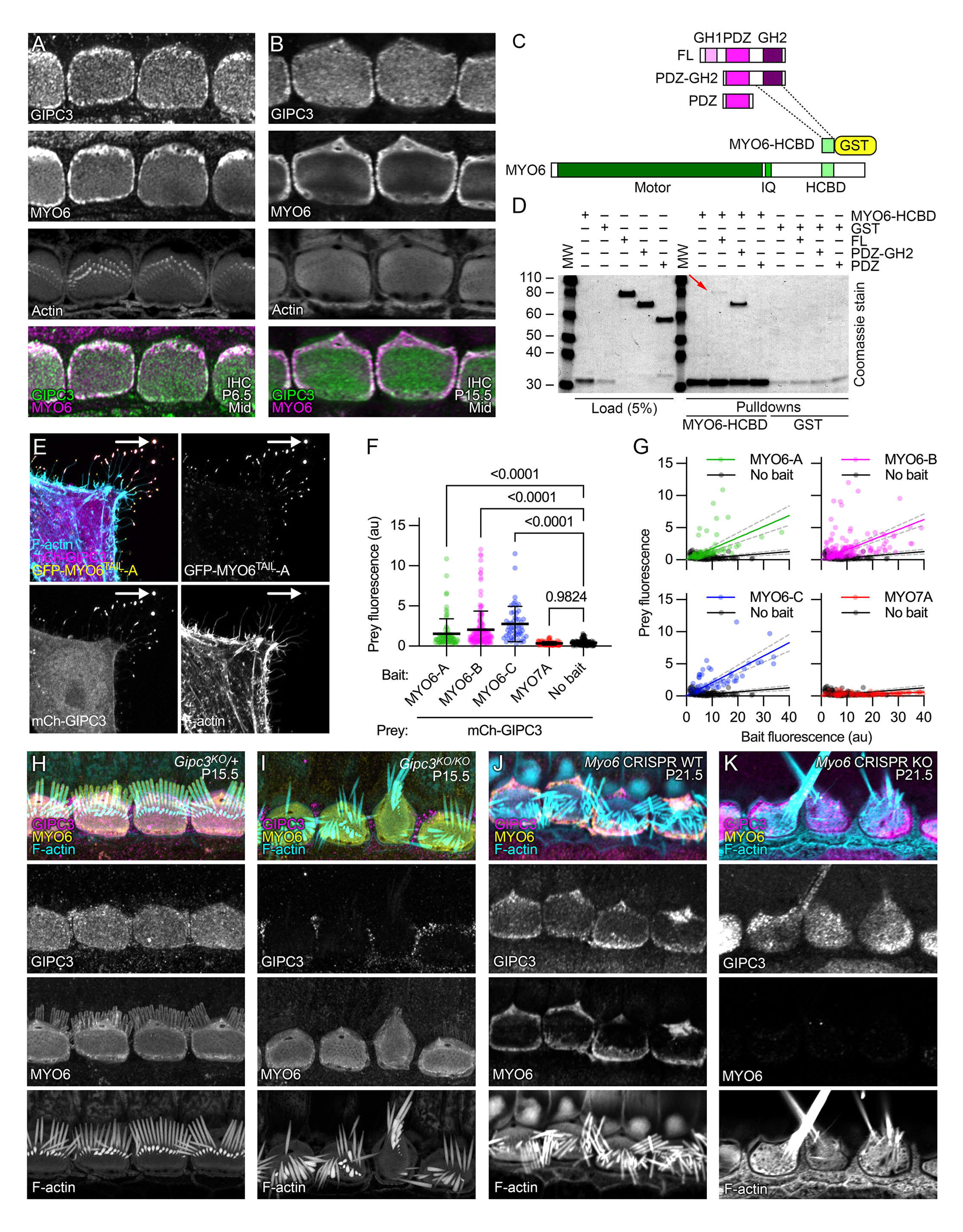
GIPC3 interacts with MYO6. ***A-B***, Colocalization of GIPC3 and MYO6 in IHCs at P6.5 and P15.5 in C57BL/6 cochleas. Localization was performed >3 times. ***C***, Domain and construct structure of GIPC3 and MYO6. ***D***, Coomassie-stained gel showing interaction of GIPC3 and MYO6 using GST pulldowns. The GIPC3 full-length construct (red arrow) and the GIPC3 construct containing the PDZ and GH2 domains both interacted with the MYO6 HCBD construct; the GIPC3 construct containing only the PDZ domain did not. Experiment was performed twice. ***E-G***, NanoSPD analysis of interactions with GIPC3. All transfected cells had mCherry-GIPC3 and MYO10^NANOTRAP^; GFP-MYO6^TAIL^ constructs were also included, which interacted with MYO10^NANOTRAP^ and were targeted to filopodial tips. ***E***, Example of interaction of mCherry-GIPC3 with GFP-MYO6^TAIL^-A. ***F***, Prey fluorescence with various constructs. Mean ± s.d. plotted. One-way ANOVA comparisons to no-bait condition with Dunnett correction for multiple comparisons. ***G***, Intensity correlation analysis, using scatter plot of bait (x-axis) and prey (y-axis) fluorescence at individual filopodia tips (from three independent determinations). We used linear fits through the origin; values for slope and R^2^ were: MYO6A-A (0.172, 0.19), MYO6A-B (0.156, 0.21), MYO6A-C (0.207, 0.39), MYO7A (0.015, -0.32), no bait (0.032, -0.85). Dashed lines are 95% confidence intervals. Sample sizes (n) were MYO6A-A (79 filopodia), MYO6A-B (160), MYO6A-C (52), MYO7A (101), no bait (116). ***H-I***, MYO6 localization did not change in *Gipc3^KO/KO^* IHCs. ***J-K***, GIPC3 was mislocalized in *Myo6* CRISPR knockout G0 IHCs. Panel widths: A-B, 25 µm; E, 30 µm; H-K, 35 µm.

We used glutathione-S-transferase (GST) pulldown experiments to confirm that GIPC3 and MYO6 interact directly. GIPC1 interacts with the helical cargo-binding domain (HCBD) of MYO6 (Shang et al., 2017), so we used either HCBD-GST or GST alone to precipitate HA-tagged GIPC3 or GIPC3 fragments containing the GIPC3 PDZ and GH2 domains or the PDZ domain alone (Fig. 3C-D). HCBD-GST precipitated full-length GIPC3, albeit only a small amount (red arrow in Fig. 3D); because both GH1 and GH2 domains are present, this molecule is likely in the autoinhibited state, incapable of binding efficiently to MYO6 (Shang et al., 2017). Deletion of the GH1 domain allowed much more strong binding to HCBD-GST, but eliminating the GH2 domain abolished the MYO6 interaction (Fig. 3D). We conclude that the GH2 domain of GIPC3 interacts with HCBD of MYO6, especially when autoinhibition is relieved.

We exploited the NanoSPD method (Bird et al., 2017) to provide additional evidence for the GIPC3-MYO6 interaction (Fig. 3E-G). In these experiments, we expressed MYO10^NANOTRAP^, a construct that targets filopodia tips of HeLa cells because of its MYO10 motor and binds to GFP fusion proteins through its nanotrap-anti-GFP single-chain antibody (Bird et al., 2017). Constructs used in NanoSPD experiments are listed in Table S1. Co-expressed GFP-tagged constructs, like the MYO6 tails, bound to MYO10^NANOTRAP^ and were also transported to filopodial tips. Proteins tagged with a fluorescent protein that does not bind to MYO10^NANOTRAP^, for example mCherry, can then be co-transported to filopodial tips if the GFP-tagged and mCherry-tagged proteins interact. As a positive control, we confirmed that when MYO10^NANOTRAP^ was co-expressed, GFP-MYO7A^TAIL^ enabled transport of mCherry-PDZD7 to filopodial tips (Fig. S2A-B) as previously reported (Morgan et al., 2016).

We compared the tail domains from three splice forms of MYO6 that were predicted in Ensembl (ensembl.org): *Myo6-212* (its protein product is referred to here as MYO6-A), *Myo6-201* (MYO6-B), and *Myo6-203* (MYO6-C); these three splice forms respectively correspond to the large-insert, small-insert, and no-insert splice forms previously reported (Buss et al., 2001). The two splice sites used for these isoforms flank the HCBD domain, raising the possibility that splicing regulates GIPC3-MYO6 interaction. Nevertheless, when co-expressed with MYO10^NANOTRAP^, each of the three splice forms of GFP-MYO6^TAIL^ transported mCherry-GIPC3 to filopodial tips of HeLa cells (Fig. 3E-G; Fig. S2C-E). mCherry-GIPC3 was located in the cytoplasm of HeLa cells when expressed alone (Fig. S2I). When expressed with MYO10^NANOTRAP^, neither GFP-MYO7A^TAIL^ (Fig. S2F) nor GFP alone (Fig. S2G) facilitated targeting of mCherry-GIPC3 to filopodial tips.

The elevation of GFP-MYO6^TAIL^ signal at filopodial tips was significant for each of the three MYO6 constructs compared to the controls (Fig. 3F). For GFP-MYO6^TAIL^ (bait) plus mCherry-GIPC3 (prey) experiments, increased prey fluorescence at filopodial tips correlated with higher levels of bait fluorescence (Fig. 3G); there was no such correlation for GFP-MYO7A^TAIL^ (Fig. 3G).

### GIPC3 is mislocalized in *Myo6* null mice

In *Gipc3^KO/KO^* mice, the distribution of MYO6 was not different from that in heterozygous controls (Fig. 3H-I); MYO6 thus did not depend on GIPC3 for localization. To test whether GIPC3 localization depends on MYO6, we used improved-genome editing via oviductal nucleic acids delivery (i-GONAD) (Ohtsuka et al., 2018), a CRISPR-Cas9 strategy, to create G0 animals that had null mutations in both *Myo6* alleles. We found a high correlation between biallelic targeting in genotyping assays of mouse tails and the loss of MYO6 immunoreactivity in hair cells (Table S2), suggesting that CRISPR-Cas9 gene modification occurred early during development. Animals with disrupted *Myo6* alleles had severely disrupted hair bundles, including fused stereocilia, as has been reported for the *Myo6^sv^* allele (Self et al., 1999).

Hair cells from i-GONAD-generated G0 mouse pups that had two wild-type *Myo6* alleles had normal distribution of GIPC3 (Fig. 3J). By contrast, in G0 pups with presumptive null *Myo6* mutations in both alleles, GIPC3 distribution was significantly perturbed (Fig. 3K); GIPC3 no longer concentrated at the hair cell periphery, and instead was found in an apparently cytoplasmic pattern. Localization of GIPC3 near cellular junctions thus depended on MYO6.

### Misshapen cuticular plates in *Gipc3^KO/KO^* inner hair cells

As IHCs of C57BL/6 mice developed, their cuticular plates elongated along the cochlear longitudinal axis (Fig. 4A). This shape transition matched that of the apical circumference, which has been described as shifting from a near-circular shape around P1 to a rounded rectangular one from P5 on (Etournay et al., 2010). While the cross-sectional area of the cuticular plate did not change between P0 and P20 (Figs. 4A and 4P, left), the vertical depth increased (Fig. 4G, Q). Using the simplifying geometric assumption that the cuticular plate was a hemisphere, we estimated that the cuticular plate volume increased ∼30% over this period (Fig. 4R, left). An alternative simplifying assumption, that the cuticular plate was a flat cylinder, gave similar results for volume.

**Figure 4.**
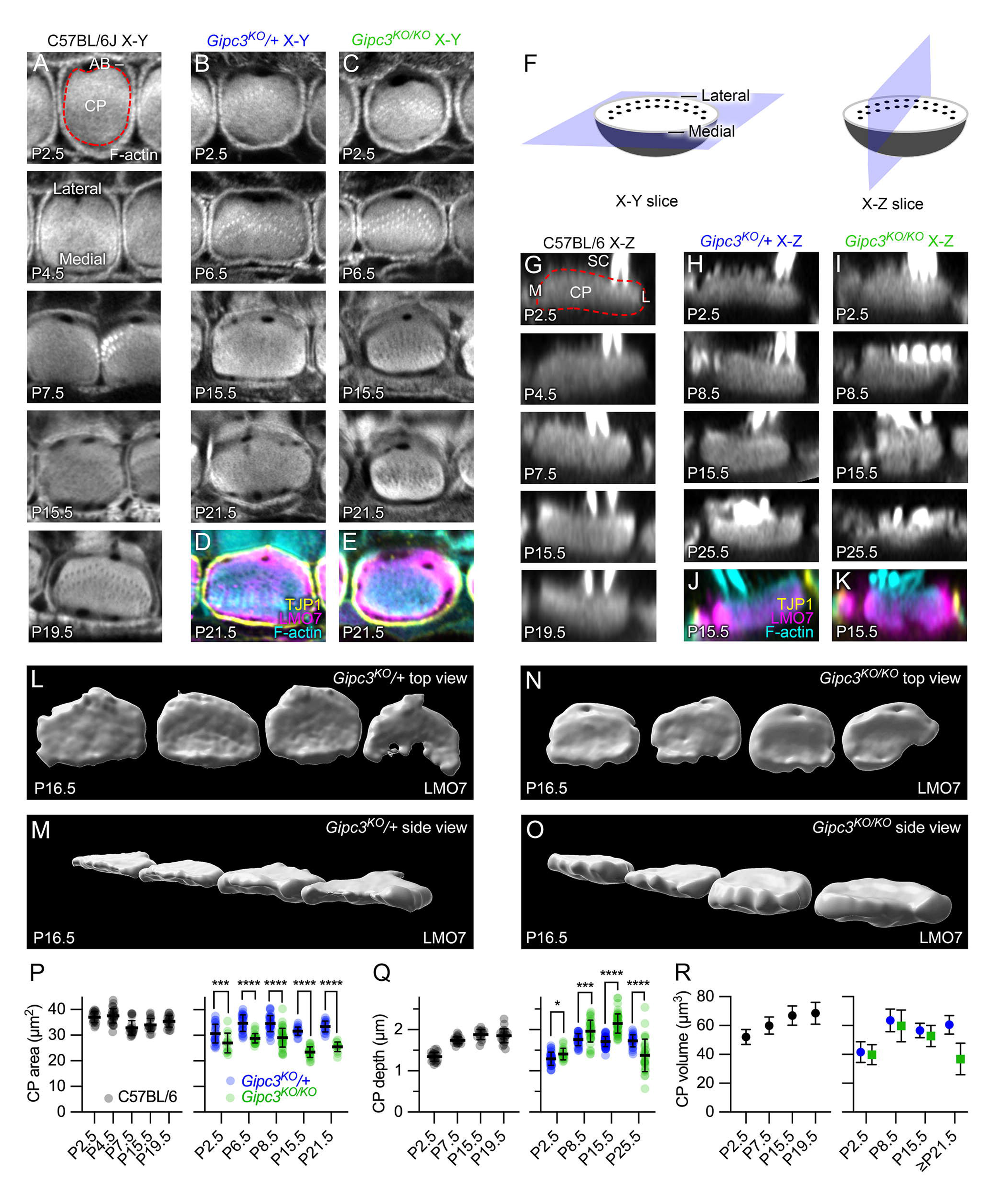
Cuticular plate defects in *Gipc3* knockout mice. ***A***, Examples of phalloidin labeling at the level of the cuticular plate for C57BL/6 IHCs (x-y slices). Red dashed line in P2.5 example outlines cuticular plate. Lateral and medial sides of the hair cell indicated in P4.5 panel. Key: CP, cuticular plate; AB, circumferential actin belt. ***B-C***, Examples of phalloidin labeling at the level of the cuticular plate for *Gipc3^KO^* heterozygote (B) and homozygote (C) IHCs. ***D-E***, Triple labeling for TJP1 (ZO1; showing apical cell junctions), F-actin, and LMO7 (showing cuticular plate). ***F***, Diagrams illustrating X-Y slice used for panels A-E and X-Z reslice used for panels G-K. Lateral and medial sides of the hair cell indicated. ***G***, Examples of phalloidin labeling through the cuticular plate for C57BL/6 IHCs (X-Z reslices). Key: SC, stereocilia; M, medial edge of hair cell; L, lateral edge. ***H-I***, Examples of phalloidin labeling through the cuticular plate for *Gipc3^KO^* heterozygote (H) and homozygote (I) IHCs. ***J-K***, Triple labeling for TJP1, actin, and LMO7. ***L-O,*** Imaris reconstruction (rendering) of LMO7 labeling in *Gipc3^KO^* heterozygote (L-M) and homozygote (N-O) IHCs at P16.5. The lateral (kinocilium) edge of the hair cell is at top in L and N; the medial edge is at the bottom. M and O are the same cuticular plates as in L and N but are rotated in two axes. ***P-R***, Quantitation of cuticular plate dimensions (mean ± s.d.). ***P***, Quantitation of IHC cuticular plate area from P2.5 to P19.5 in C57BL/6 IHCs (left) and from P2.5 to P21.5 *Gipc3^KO^* heterozygote and homozygote IHCs (right). For *Gipc3^KO^*, unpaired t-tests were used for statistical comparisons. Mean ± s.e.m. are plotted in right panels (also for M and N). P values for cuticular plate area were: P2.5, 0.0073 (n=24 and 13); P6.5, <0.00001 (n=28 and 27); P8.5, <0.00001 (n=32 and 41); P15.5, <0.00001 (n=18 and 25); P21.5, <0.00001 (n=17 and 14). ***Q***, Quantitation of cuticular plate depth from P2.5 to P19.5 in C57BL/6 IHCs (left) and from P2.5 to P25.5 in *Gipc3^KO^* heterozygote and homozygote IHCs (right). P values for cuticular plate depth were: P2.5, 0.011 (n=33 and 19 for heterozygote and knockout); P8.5, 0.0003 (n=30 and 35); P15.5, <0.0001 (n=60 and 53); P21.5, <0.0001 (n=27 and 31). ***R***, Quantitation of cuticular plate volume from P2.5 to P19.5 in C57BL/6 IHCs (left) and from P2.5 to ≥P21.5 in *Gipc3^KO^* heterozygote and homozygote IHCs (right). Panel widths: A, 17.5 µm; B-K, 12 µm; L-O, 50 µm.

In *Gipc3^KO/KO^* IHCs, the shift of the cuticular plate circumference from circular to rounded rectangular did not occur (Fig. 4B-C). The cross-sectional area of the *Gipc3^KO/KO^* cuticular plate was reduced compared to *Gipc3^KO^*/+ at all time points during development (Fig. 4P, right). The decrease in cuticular-plate area was initially offset by a corresponding increase in cuticular plate depth (Fig. 4H-I, Q), however, such that the estimated cuticular-plate volume in *Gipc3^KO^*/+ and *Gipc3^KO/KO^* IHCs was nearly identical (Fig. 4R, right). After P21, the cuticular plate thinned considerably in *Gipc3^KO/KO^* hair cells (Fig. 4Q, right), which led to a decrease in cuticular plate volume (Fig. 4R, right). GIPC3 thus plays a role in the developmental elongation and flattening of IHC cuticular plates.

Labeling for components of the apical region highlighted these changes. We used antibodies against TJP1 (ZO1) to label apical junctions and antibodies against LMO7 to label the cuticular plate (Du et al., 2019). In *Gipc3^KO/KO^* mutants, LMO7 labeling remained throughout the apical area even though the cross-sectional area of the cuticular plate, as determined by phalloidin labeling, decreased (Fig. 4D-E).

We used image rendering to reconstruct cuticular plates marked by LMO7 labeling at P16.5 (Fig. 4L-O). The distinction between the thin, flattened cuticular plates of *Gipc3^KO^*/+ controls (Fig. 4L-M) and the more rounded cuticular plates of *Gipc3^KO/KO^* mutants (Fig. 4N-O) was apparent. Volumes were directly estimated from the rendered LMO7 images, not using simplifying geometric assumptions; *Gipc3^KO^*/+ control LMO7 volumes were 63 ± 18 µm^3^ (mean ± s.d.; n=13), while *Gipc3^KO/KO^* mutants were 71 ± 18 µm^3^ (*P* = 0.165; Student’s t-test).

### Altered apical junctions of *Gipc3^KO/KO^* hair cells

IHCs are packed closely together along the cochlear axis, albeit separated by luminal processes of inner phalangeal cells (IPhCs) (Driver and Kelley, 2009). In control IHCs, antibodies against TJP1 labeled the junctional regions of IHCs, but not the IPhCs, revealing two parallel lines of immunostaining separated by a very small gap (Fig. 5A, arrows). Examination of a serial block-face scanning electron microscopy dataset from mouse cochlea (Hua et al., 2021) showed that two adjacent IHCs sandwiched microvilli projecting from the IPhCs; the microvilli and IHC plasma membranes appeared to be close enough that the interaction was mediated by cell-cell contacts (Fig. S3). In addition, some direct IHC-IHC contacts also appeared to be present in areas with fewer microvilli (Fig. S3).

**Figure 5.**
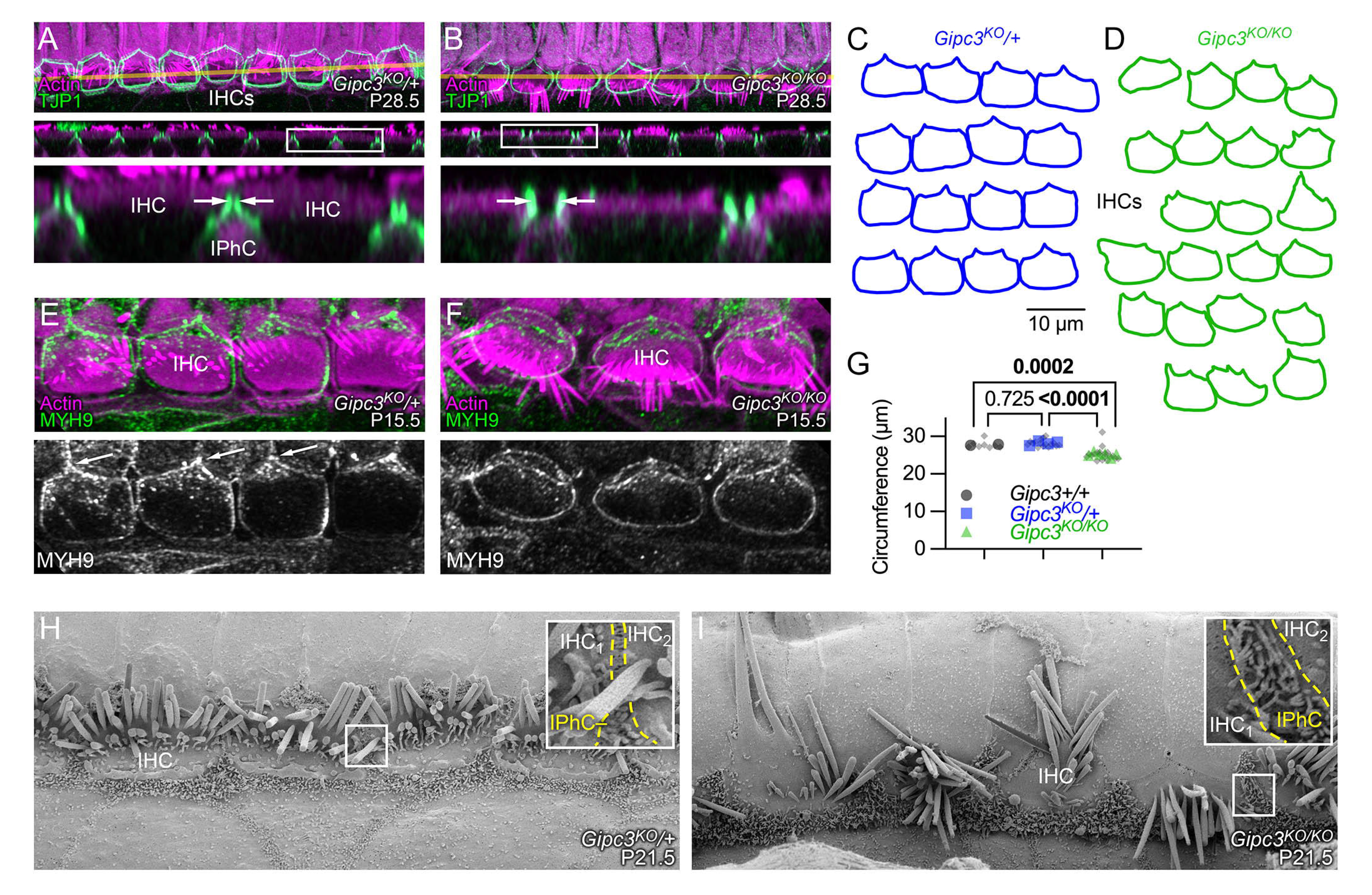
Apical junction defects in *Gipc3* knockout mice. ***A-B***, TJP1 immunoreactivity highlights apical junctions in *Gipc3^KO^* IHCs. X-Y (top) and X-Z (middle) slices. Yellow lines in upper panels indicate transects used for X-Z reslices in middle panels; magnified boxed regions of reslices are below. immunolocalization experiments were performed >3 times. Key: IPhC, inner phalangeal cell. ***C-D***, Tracings of TJP1 labeling from four *Gipc3^KO^*/+ (C) and six *Gipc3^KO/KO^* (D) cochleas. ***E-F***, MYH9 immunoreactivity highlights apical junctions in *Gipc3^KO^* cochleas. ***G***, Apical circumference area for the indicated genotypes. *P*-values determined from nested one-way ANOVA with Tukey correction. ***H-I***, Scanning electron microscopy of P21.5 *Gipc3^KO^*/+ and *Gipc3^KO/KO^* IHC region. Insets show the junction region between an IHC, an IPhC, and the next IHC. The yellow dashed lines outline the region in between the two IHCs that is occupied by IPhC microvilli. P21.5 SEM was carried out once, with similar results seen in >25 IHCs in each genotype. Panel widths: A-B upper and middle panels, 80 µm; A-B lower panels, 20 µm; E-F, 35 µm; G-H, 30 µm (insets, 2 µm).

In *Gipc3^KO/KO^* cochleas, however, the gap between IHCs widened noticeably (Fig. 5B, arrows); this space was presumably filled in by expansion of the apical surfaces of IPhCs. The consistent difference between the association of adjacent IHCs is illustrated in Fig. 5C-D; while *Gipc3^KO^*/+ IHC outlines were relatively rectangular, *Gipc3^KO/KO^* IHC outlines were considerably more rounded and did not appear to be tightly coupled together. In addition, the apical circumference of *Gipc3^KO/KO^* IHCs was reduced significantly when compared to wild-type or heterozygote IHCs (Fig. 5G).

Immunoreactivity for the MYH9 (nonmuscle myosin IIa) in *Gipc3^KO^*/+ and *Gipc3^KO/KO^* IHCs (Fig. 5E-F) highlighted the relatively square appearance of the apical junction region in *Gipc3^KO^*/+ compared to the more rounded appearance in *Gipc3^KO/KO^*. In *Gipc3^KO^*/+, MYH9 was concentrated at the junction between one IHC and two pillar cells, a tripartite junction that forms an apex pointing laterally in the cochlea (Fig. 5E, arrows). By contrast, MYH9 was relatively uniformly distributed around the apical junctional region in *Gipc3^KO/KO^* IHCs (Fig. 5F).

The loss of close opposition of IHCs in *Gipc3^KO/KO^* cochleas was apparent by SEM (Fig. 5H-I). In heterozygotes, a few IPhC microvilli projected between the apical surfaces of two adjacent IHCs, which appeared to have some direct contacts (Fig. 5H inset). By contrast, the microvilli-endowed apical surfaces of IPhCs expanded substantially in *Gipc3^KO/KO^* cochleas, and apical surfaces of IHCs flanking each IPhC were relatively far apart (Fig. 5I inset).

### Immunoaffinity purification of GIPC3 protein complexes

To understand how GIPC3 exerts its effects on dimensions of cuticular plates and apical surfaces, we examined the GIPC3 protein-interaction network in hair cells. Because of its superior recognition of chicken GIPC3, we exploited the 10G5 anti-GIPC3 monoclonal antibody to immunoaffinity purify GIPC3 from crosslinked chicken inner ear extracts. We used a fraction that was enriched for stereocilia, but still contained large amounts of hair-cell cytoplasmic proteins (Morgan et al., 2016). We carried out two separate experiments, each with ∼1000 chicken ears, where we stabilized protein complexes using primary amine-reactive homo-bifunctional N-hydroxysuccimide ester crosslinkers that are thiol-cleavable and hence reversible (Mattson et al., 1993). In one experiment, we used dithiobis(succinimidyl propionate) (DSP), a membrane-permeable crosslinker that crosslinks extracellular and intracellular complexes; in the other experiment, we used 3,3’-dithiobis(sulfosuccinimidyl propionate) (DTSSP), which is membrane impermeant and thus only stabilizes extracellular and transmembrane complexes. We prepared soluble extracts of crude, crosslinked stereocilia (S7 in Fig. 6A) and used these fractions (S7 from DSP or DTSSP) for identifying GIPC3-interacting proteins.

**Figure 6.**
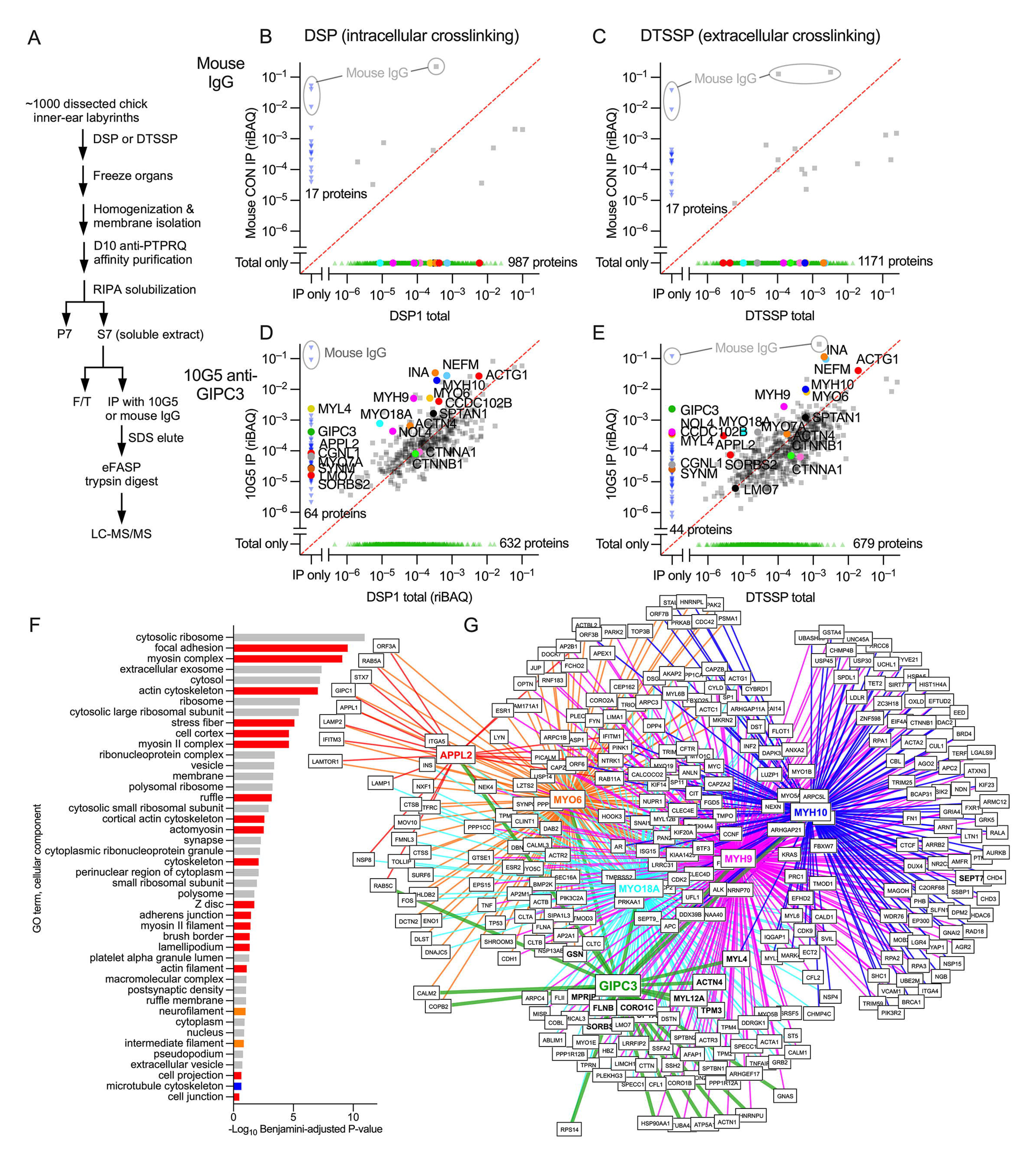
GIPC3 interaction networks identified through immunoaffinity purification and protein mass spectrometry. ***A***, Flow chart for anti-GIPC3 immunoaffinity purification from crude chick stereocilia extracts. ***B-E***, Comparison of abundance (riBAQ) of proteins detected in DSP1 total or DTSSP total (starting S7 extract; plotted on x-axis) compared to the immunoprecipitates (y-axis) for mouse IgG control (B-C) and 10G5 anti-GIPC3 (D-E) experiments. Panels B and D show results with the DSP-crosslinked starting extract, while panels C and E show results with DTSSP crosslinking. Each point represents that protein’s average abundance in two experiments (biological replicates); symbol colors are arbitrarily chosen. Red dashed line is the unity line (equal riBAQ in total and IP). Key proteins are called out. Mouse IgG protein from immunoprecipitation is highlighted in gray. ***F***, Gene ontology analysis (cellular component) with DAVID of the top 50 proteins from the DTSSP 10G5 eluate. Color code: Red, actin-associated components; orange, intermediate filament-associated components; blue, microtubule-associated components; gray, other components. ***G***, Overlap of protein interaction networks of key proteins from the DTSSP 10G5 eluate (GIPC3, APPL2, MYO6, MYO18A, MYH9, and MYH10). BioGRID-defined protein networks for APPL2, MYO6, MYO18A, MYH9, and MYH10 were compared with the top 100 proteins from the DTSSP 10G5 eluate. Only proteins identified as interactors of two or more of the key proteins were included. Proteins in bold were present in the top 100 proteins from the DTSSP 10G5 eluate.

When proteins were crosslinked with DSP, very few proteins were purified from S7 by control mouse IgG (Fig. 6B). By contrast, 429 proteins were precipitated from S7 (out of 1061 total) with 10G5 (Fig. 6D). Many proteins were enriched substantially by precipitation—their relative abundance in the immunoaffinity precipitate was greater than that in the starting material—including GIPC3 itself, which was detected in the 10G5 immunoaffinity precipitate but not in the S7 starting extract. The immunoaffinity purification was repeated using the flow-through from this DSP extract experiment as the starting material, which gave very similar results (Fig. S4), which suggested that the antibody-coupled beads were saturated with GIPC3 and its partners in the first immunoprecipitation. Immunoaffinity purification results with DTSSP were broadly similar, although GIPC3 made up a larger fraction of the precipitate and other proteins were at lower levels in the precipitate than in the DSP experiment (Fig. 6D, E), consistent with the lack of intracellular crosslinking.

Gene ontology (GO) analysis of the top 35 proteins most enriched in the GIPC immunoprecipitations (Table S3) indicated that many were associated with actin components (red in Fig. 6F). Key cellular component terms included focal adhesion, myosin complex, actin cytoskeleton, stress fiber, cell cortex, and myosin II complex. This analysis suggested that the GIPC3 complexes purified included co-precipitating fragments of adherens junctions, circumferential actin belts, or cuticular plates. These results are consistent with localization of GIPC3 in these regions and morphological disruptions to the same areas in the *Gipc3^KO/KO^* mice.

To determine how the protein networks of key immunoprecipitated proteins overlapped, we examined pairwise the binary interactions listed in the BioGRID protein-interaction database (Oughtred et al., 2021) for APPL2, MYO6, MYO18A, MYH9, and MYH10. We chose these proteins based on their known interaction with actin structures at cell junctions (MYO18A, MYH9, and MYH10) or their known interaction with the paralog GIPC1 and association with early endosomes (APPL2 and MYO6). We compared the overlap of these six networks (including GIPC3), identifying interacting proteins shared by two or more networks. Many interactions were shared by these six proteins (Fig. 6G), including many proteins that were identified in the immunoprecipitation experiments (bold in Fig. 6G). Color coding was the same as in Fig. 6B-E. The APPL2 network overlap was sparser than those of the other proteins, but still many proteins were shared with the other target proteins.

### Binding partners for GIPC3 are located in hair cells

As GIPC3 is likely to interact with target proteins using its PDZ domain, we inspected the top 35 enriched proteins for C-terminal PBMs (Table S3). Four proteins had PBMs that met the consensus for high-affinity binding to the PDZ domains of the GIPC family (Table S4): APPL2, MYO18A, ACTN1, and ACTN4 (Fig. 7A).

**Figure 7.**
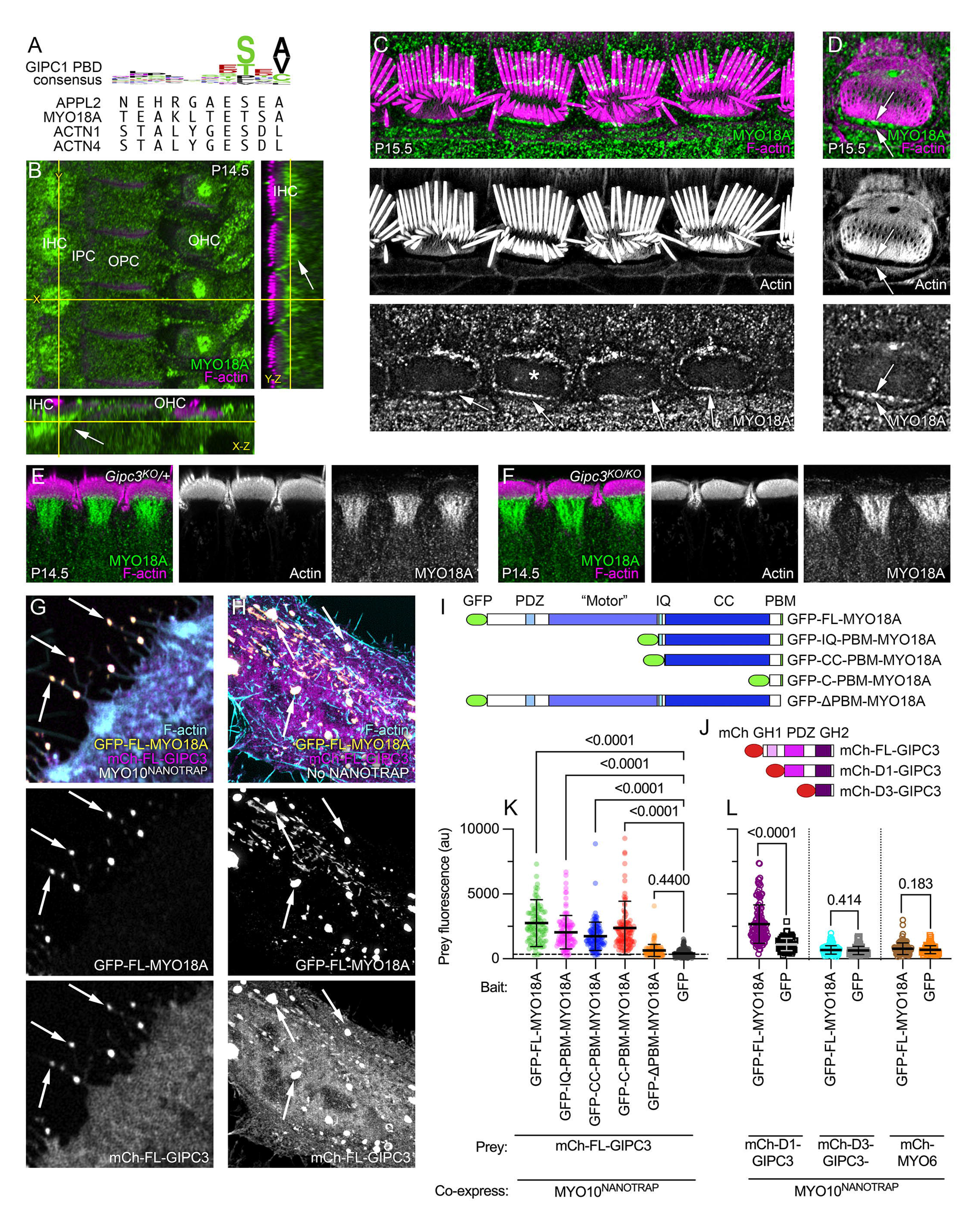
MYO18A is located in the hair cell apical domain. ***A***, Top, sequence logo for binding of ligands to GIPC1 PDZ domain; bottom, C-terminal ten amino acids of APPL2, MYO18A, ACTN1, and ACTN4. ***B***, Immunolocalization of MYO18A in P14.5 mouse cochlea; slices from a three-dimensional image stack. Transects for other image axes are shown in yellow; the X and Y transects in the main X-Y image show the locations for the Y-Z and X-Z images. Arrow indicates concentration of MYO18A immunoreactivity below the IHC cuticular plate. Key: IHC, inner hair cell; IPC, inner pillar cell; OPC, outer pillar cell; OHC, outer hair cell. immunolocalization experiments for MYO18A were performed >5 times. ***C-D***, MYO18A immunoreactivity in P15.5 IHCs using lattice SIM imaging. ***C***, Image showing four IHCs at the stereocilia/cuticular plate level. ***D,*** Image showing a single IHC (labeled with asterisk in C) at the cuticular plate level (different plane than in C). Arrows delineate the gap between cuticular plate actin and the circumferential actin belt. ***E-F***, MYO18A immunoreactivity in P14.5 IHCs from folded cochleas using Airyscan imaging. E is from a *Gipc3^KO^*/+ mouse and F is from a *Gipc3^KO/KO^* mouse. ***G***, NanoSPD of MYO18A-GIPC3. Example of filopodial targeting of mCherry-GIPC3 by GFP-FL-MYO18A, mediated by MYO10^NANOTRAP^. ***H***, Expression of FL-GFP-MYO18A and FL-mCherry-GIPC3 constructs in HeLa cells (no MYO10^NANOTRAP^ expressed). Arrows indicate large cytoplasmic aggregates containing GFP and mCherry. ***I***, GFP-MYO18A constructs. Key: “Motor,” actin- and ATP-binding domains homologous to myosin motor domains in active myosins; IQ, isoleucine/glutamine calmodulin-binding; PBM, PDZ-binding motif. ***J***, mCherry-GIPC3 constructs. Key: mCh, mCherry; GH1, GIPC-homology 1; GH2, GIPC-homology 2. ***K***, Prey (mCh-GIPC3) fluorescence with GFP-MYO18A constructs or GFP control. Mean ± s.d. plotted in K and L. One-way ANOVA comparisons to no-bait condition with Dunnett correction for multiple comparisons. Sample sizes (n) were GFP-FL-MYO18A (105 filopodia), GFP-IQ-PBM-MYO18A (109), GFP-CC-PBM-MYO18A (116), GFP-C-PBM-MYO18A (129), GFP-ΔPBM-MYO18A (127), GFP (208). ***L***, Prey (mCh-GIPC3 constructs or mCh-MYO6) fluorescence with GFP-MYO18A constructs or GFP control. One-way ANOVA comparisons to no-bait condition with Dunnett correction for multiple comparisons. Sample sizes (n) were mCh-D1-GIPC3 (128 filopodia for GFP-FL-MYO18A and 88 for GFP alone), mCh-D3-GIPC3 (180 and 47), and mCh-MYO6 (107 and 93). Panel widths: B, 37.5 µm for X-Y plot (same scale applies to Y-Z and X-Z panels); C, 50 µm; D, 12 µm; E-F, 45 µm; G-H, 15 µm.

APPL2’s paralog APPL1 is a well-characterized partner of GIPC1 (Lin et al., 2006, Varsano et al., 2006), suggesting that a GIPC3-APPL2 interaction is plausible. Indeed, we found using NanoSPD that mCherry-GIPC3 interacts with GFP-APPL2 (Fig. S2M). Immunoreactivity for APPL2 was detected in inner pillar cells (Fig. S5); although APPL2 was not definitively located in IHCs, *Appl2* transcripts were enriched in cochlear hair cells (umgear.org). These results suggest that our antibody is insufficiently sensitive or that its epitope is masked in hair cells.

*Myo18a* transcripts were highly enriched in hair cells (umgear.org) and MYO18A protein was enriched in cochlear and vestibular hair cells (Fig. 1A-B), supporting MYO18A as a candidate for interaction with GIPC3. Using the Atlas antibody specific for MYO18A (Fig. S6A-B), we found in C57BL/6 and *Gipc3^KO^*/+ heterozygote IHCs that MYO18A immunoreactivity surrounded the cuticular plate, especially underneath it (Fig. 7B). Similar results were seen with a second antibody (Fig. S5D). Lattice SIM revealed significant MYO18A immunoreactivity at the apical periphery of IHCs (Fig. 7C-D), similar to the location of GIPC3. MYO18A immunoreactivity was similarly distributed in *Gipc3^KO/KO^* mutants and heterozygote controls (Fig. 7E-F).

Alpha-actinins have been localized to hair-cell cuticular plates (Slepecky and Chamberlain, 1985). *Actn1* and *Actn4* transcripts were present in cochlear hair cells, although not enriched (umgear.org). We detected a modest increase of ACTN4 labeling in IHC cuticular plates compared to cell bodies; expression levels were much higher in pillar cells (Fig. S5C). We were unable to detect ACTN1.

Other proteins may bind to GIPC3 in hair cells. Although MYO6 is responsible for strong adhesion of cell-cell contacts mediated by CDH1 (E-cadherin) (Maddugoda et al., 2007), CDH1 expression was low in IHCs (Etournay et al., 2010). CDH2 has reciprocal expression in the cochlea, however, and was high in IHCs (Etournay et al., 2010, Simonneau et al., 2003), raising the possibility that MYO6 plays a role in CDH2-mediated cell-cell contacts. CTNNB1 (beta catenin) binds directly to CDH1 and CDH2 (Valenta et al., 2012); CTNNB1 and its partner CTNNA1 were co-purified with GIPC3 in immunoaffinity purification experiments (Fig. 6D-E), albeit not enriched relative to the starting extract. CTNNB1 has a C-terminal PBM (NQLAWFDTDL) that is similar to the GIPC3 consensus sequence; in NanoSPD experiments, we demonstrated that mEMERALD-CTNNB1 interacted with mCherry-GIPC3 (Fig. S2O-P). CTNNB1 could thus mediate interaction of GIPC3 with adhesion complexes, presumably through CTNNB1’s PBM.

### MYO18A forms aggregates and interacts with GIPC3

Experiments using NanoSPD demonstrated that full-length MYO18A tagged at the N-terminus with GFP interacted with GIPC3 (Fig. 7G). A comparison of MYO18A deletion constructs (Fig. 7I) showed that constructs that sequentially lacked the N-terminal extension, motor domain, IQ calmodulin-binding domain, and coiled-coil domain still interacted with GIPC3; only when the C-terminal PBM was deleted was the interaction of MYO18A and GIPC3 significantly reduced (Fig. 7K; Fig. S6-S7). Unsurprisingly, the interaction of MYO18A with GIPC3 required the presence of the GIPC3’s PDZ domain (Fig. 7J, L; Fig. S6-S7). MYO6 did not interact with MYO18A (Fig. 7L).

When expressed as a full-length protein without MYO10^NANOTRAP^ co-expression, MYO18A formed large aggregates within the cytoplasm of HeLa cells (Figs. 7H, S6D), resembling biomolecular condensates arising from self-association (Banani et al., 2017). These aggregates were still seen with constructs that lacked the N-terminal extension, motor domain, IQ calmodulin-binding domain, or C-terminal PDB (Fig. S6C-G). A construct with the C-terminus alone, including the PBM, was cytoplasmic (Fig. S6H), however, implicating the coiled-coil domain in mediating MYO18A self-association. Formation of aggregates suggested that at least two MYO18A-MYO18A interaction domains exist within the coiled-coil domain.

When co-expressed with GFP alone, mCherry-GIPC3 was largely cytoplasmic, although some mCherry-GIPC3 did target to an unknown target in the HeLa cell cytoplasm (Fig. S6I). By contrast, most mCherry-GIPC3 was recruited to aggregates when GFP-FL-MYO18A was co-expressed (Fig. S6J), which indicates that GIPC3 does not obscure the domains responsible for aggregation. Removal of the N-terminal extension, motor domain, IQ calmodulin-binding domain, or coiled-coil domain had no effect on recruitment of mCherry-GIPC3 to GFP-MYO18A aggregates (Fig. S6K-L). While aggregates were absent when C-PBM-MYO18A was expressed, localization of co-expressed mCherry-GIPC3 still matched that of C-PBM-MYO18A (Fig. S6M). By contrast, when the four C-terminal amino acids of FL-MYO18A were deleted, mCherry-GIPC3 no longer interacted with MYO18A (Fig. S6N). Together the experiments of Figs. 7 and S6 demonstrate that the PBM of MYO18A interacts with the PDZ domain of GIPC3.

## Discussion

Our data show that GIPC3 is essential for hair-cell function, and roles for this protein include shaping the cuticular plate and contributing to normal cell-cell junctions. Our results confirm that the GH2 domain of GIPC3 interacts with the molecular motor MYO6; moreover, we show that the PDZ domain interacts with a number of proteins situated to be involved in cuticular-plate and junction function, including MYO18A and the α-actinins ACTN1 and ACTN4. Moreover, the GIPC3 network also includes the nonmuscle myosin II proteins MYH9 and MYH10, perhaps coupled through their binding partner MYO18A (Billington et al., 2015). Our working hypothesis is that GIPC3 establishes multiple complexes that include MYO6 and other key molecules, and that these complexes operate at the apex of the hair cell to connect the cuticular plate to apical junctions, directly or indirectly.

### Cuticular plate and hair-bundle shape changes in *Gipc3^KO/KO^*

Cochlear hair cells undergo a notable change in shape of their apical region in the early postnatal period (Fig. 8A), which has been ascribed to internal tension generated by MYO7A (Etournay et al., 2010). Disruption of hair-bundle integrity interferes with the apical shape change, which suggests a reciprocal interaction between stereocilia and their rootlets with the cuticular plate and apical junction complex (Etournay et al., 2010). GIPC3-engaged MYO6 may also contribute to this internal tension, albeit with a less dramatic impact on bundle integrity and without the loss of MET current.

**Figure 8.**
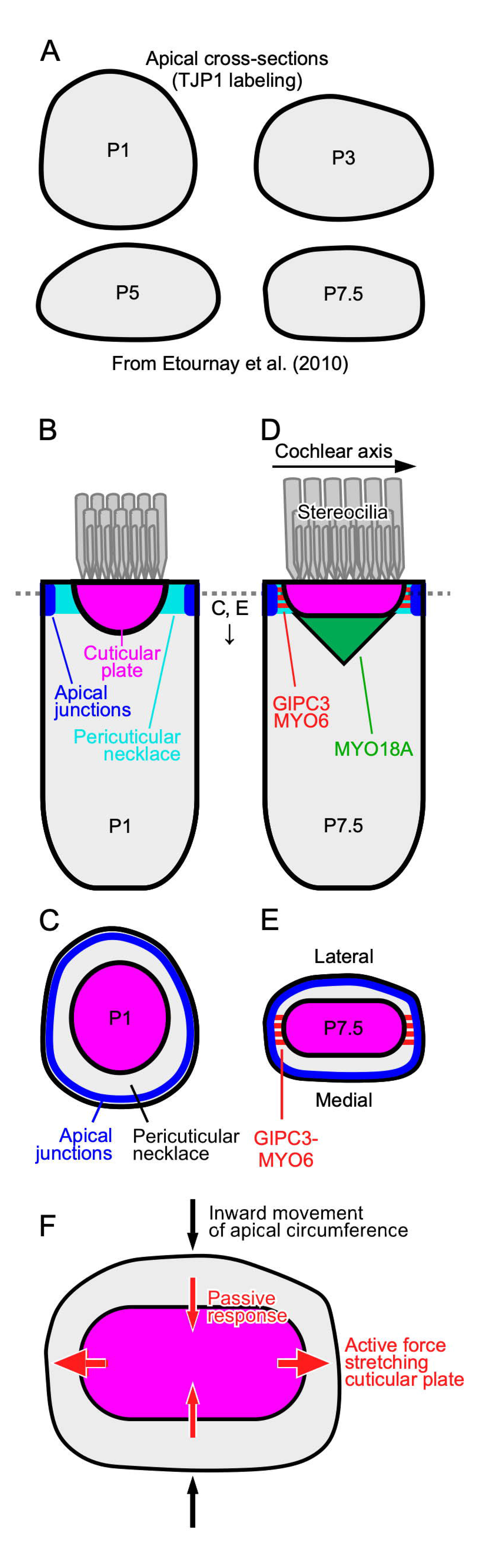
Model for GIPC3 coupling of apical cell junctions to the cuticular plate. ***A***, Tracings of averaged TJP1-labeled apical cell borders at indicated ages (adapted from Fig. 2 of Etournay et al., 2018). ***B-C***, Key complexes and processes in P1 IHC. Cuticular plate is rounded up early in development. Apical junctions in blue, cuticular plate in dark orange, stereocilia in gray. The pericuticular necklace is the gap between the apical junctions and the cuticular plate. MYO18A is not shown at this age. ***D-E***, P7.5 IHC. Cuticular plate is flattened later in development. GIPC3-MYO6 complexes in red; MYO18A structure in green; MYO18A at apical junction region is not shown. ***F***, As the apical circumference is remodeled between P1 and P7.5, the IHC narrows along the lateral-medial axis. Inside the cell, active force generated along the cochlear axis stretches the cuticular plate; maintaining constant volume, the cuticular plate passively shrinks along the lateral-medial axis. GIPC3-MYO6 complexes either generate or are simply coupled to the force production that stretches the cuticular plate. Loss of GIPC3 prevents the elongation of the cuticular plate. The MYO18A structure underneath the cuticular plate may assist in its flattening; GIPC3 could couple MYO18A there with MYO6, ACTN1, or ACTN4 in the cuticular plate. B and D show medial-lateral sections through IHC centers; C and E-F show cross-section of IHCs at level of dashed line in B and D. The pericuticular necklace was removed for clarity in C-F, and the apical junctions were removed in F.

A prominent consequence of the loss of GIPC3 in hair cells was that the cuticular plate remained more round, both cell’s apical-to-basal axis (Z axis) and in the perpendicular axis (X-Y axis). In *Gipc3^KO/KO^* IHCs, the cross-sectional area of the cuticular plate never reached the size seen in *Gipc3^KO^*/+ controls, while the depth of the cuticular plate did not reduce as it does in in controls. Accordingly, the volume of the cuticular plate did not change appreciably until after P15.5 in *Gipc3^KO/KO^* IHCs, suggesting that the cuticular plate was formed normally early during development but was then subjected to different internal forces in the mutant. In particular, the cuticular plate did not flatten and extend normally during development in *Gipc3^KO/KO^* IHCs, i.e., it did not proceed from hemisphere-like to disk-like (Fig. 8).

*Gipc3^KO/KO^* hair bundles differed from those of *Gipc3^KO^*/+ controls in several ways. Stereocilia in IHC bundles appeared to be pushed together, and shorter stereocilia were generally thicker than shorter stereocilia of controls. Maximum transduction currents were higher in *Gipc3^KO/KO^* IHCs, suggesting that there were more channels, but offset stimuli were required to fully elicit the maximum current. Both IHC and OHC bundles of *Gipc3^KO/KO^* mice had a squeezed appearance, where the flanking wings of the bundle (stereocilia most distal from the fonticulus and basal body) were closer together. This phenotype correlated well with the lack of flattening and extension of the cuticular plate, as if the stereocilia were coupled within the cuticular plate and were, similar to the cuticular plate, not subjected to the internal forces that spreads them out along the cochlear axis.

### GIPC3’s protein network

Immunoaffinity purification experiments with anti-GIPC3 precipitated many cytoskeletal and junctional proteins, presumably in large complexes that were crosslinked either both extracellularly and intracellularly (with DSP experiments) or extracellularly alone (DTSSP). The molar abundance in the immunoprecipitates (estimated from the Y-axis in Fig. 6D-E) of most cytoskeletal proteins was much greater than that of GIPC3, which suggests that a small number of GIPC3 molecules engaged large cytoskeletal networks. These networks likely included the cuticular plate itself, as well as the apical cell junctions connecting hair cells to surrounding supporting cells. In addition, cytoskeletal proteins concentrated in the pericuticular necklace were likely also present, given GIPC3’s location there.

GIPC3 includes a PDZ domain, and several proteins enriched in the GIPC3 immunoaffinity purification experiments contained sequences that matched the consensus C-terminal PBM for the GIPC family. Although these immunoaffinity purification experiments were carried out with chick inner ear, we confirmed interaction of mouse MYO18A, APPL2, and CTNNB1 with mouse GIPC3 using NanoSPD assays.

The interaction of GIPC1 and MYO6 is well established, and preliminary evidence suggested that GIPC3 and MYO6 interact (Shang et al., 2017). We confirmed this GIPC3-MYO6 interaction using in vitro GST-pulldown experiments, which showed that the GH2 domain of GIPC3 interacted with the helical cargo-binding domain (HCBD) of MYO6. Moreover, immunoaffinity purification and NanoSPD experiments showed that MYO6 associates with complexes containing GIPC3, presumably because of the direct GIPC3-MYO6 interaction. Additional evidence for this interaction includes similar phenotypes of the *Myo6* and *Gipc3* null mouse lines, as well as the mislocalization of GIPC3 in *Myo6* knockout model that was generated using i-GONAD and CRISPR/Cas9. In particular, the latter results showed that MYO6 is responsible for localization of GIPC3 at the apical periphery of hair cells, near cell-cell junctions. Full-length GIPC3 forms an autoinhibited dimer, and so the interaction with MYO6 likely occurs only after GIPC3 has been activated by a ligand with a PBM that binds the GIPC3 PDZ domain (Shang et al., 2017), which in turn suggests that GIPC3 not only interacts with MYO6 at the apical periphery, but also with an activating ligand.

These experiments highlighted MYO18A, another unconventional myosin that is expressed in hair cells. Hair cells are notable for their reliance on myosins, presumably because their actin-rich cytoskeletal structures, especially the stereocilia and cuticular plate, are substrates for myosins (Friedman et al., 2020, Hasson et al., 1997). The location of MYO18A below the cuticular plate suggests that it is involved in constraining the cuticular plate, a role also proposed for the striated organelle, an enigmatic structure also found in hair cells below the cuticular plate (Slepecky et al., 1981, Vranceanu et al., 2012). MYO18A is plausibly a component of the striated organelle. In addition, MYO18A localizes near apical junctions connecting IHCs to surrounding supporting cells, like the apical distribution of GIPC3. Mice homozygous for a global *Myo18a* null allele show preweaning lethality with complete penetrance (www.mousephenotype.org), thus preventing simple genetic analysis.

### Model for cuticular plate shaping

Etournay et al. (2010) showed that apical surfaces of IHCs shift from near circular to rounded rectangular by P7.5 (Fig. 8A); the distance along the cochlear axis remained approximately the same, however, while the distance along the lateral-medial axis shortened (Etournay et al., 2010). Our results suggest that force is generated during this shape change that is also coupled to lengthening of the cuticular plate along the same axis, and that transmission of this force requires GIPC3. As development proceeds, the apical circumference remodels, the cuticular plate flattens, and anchoring at the plate’s sides and base ensures that it forms an immovable platform for insertion of stereocilia. Loss of GIPC3 does not affect the total amount of cuticular plate material, however, as the cuticular plate’s volume was not altered in *Gipc3^KO/KO^* hair cells. As internal forces are generated, the cuticular plate extends; because a constant volume is maintained, the plate thins (Fig. 8). The hair-bundle phenotype suggests that the stereocilia are moved into place because of their anchoring within the cuticular plate; bundles start as a tight cluster of stereocilia (Kaltenbach et al., 1994), but as development proceeds, they are separated into rows and columns by the forces extending the cuticular plate. When GIPC3 is absent, this separation does not occur properly.

Based on localization of MYH9 and MYO7A, apical circumference remodeling may result from forces applied within hair cells (Etournay et al., 2010). MYO18A itself likely cannot generate this tension; it has very low myosin ATPase activity (Guzik-Lendrum et al., 2013), although it can form mixed filaments with MYH9 (Billington et al., 2015). We suggest that GIPC3-MYO6 complexes assist in anchoring the cuticular plate to apical cell-cell junctions and coupling to internal forces (Fig. 8B-E); whether MYO6 contributes to force elongating the cuticular plate or simply acts as an anchor is unknown.

Several mechanisms are plausible for formation of the rounded rectangular apical circumference in IHCs (Fig. 8). For example, internal contraction force could be generated along the lateral-medial axis, squeezing these two sides of the hair cell closer together (Fig. 8F). Given that the cell’s apical surface area does not change, tension could then develop within the cell along the cochlear axis; this tension could be used to reshape the cuticular plate if it was coupled to the apical junctions. Alternatively, forces may be applied to hair cells externally by the surrounding supporting cells; supporting cells could squeeze IHCs along their lateral-medial axis or elongate them along the cochlear axis. The rounded shape of cuticular plates in *Gipc3^KO/KO^* IHCs argues against this possibility, however, and suggests that the forces are generated internally. Whether reshaping forces are generated internally or externally, GIPC3 and MYO6 may be located in the ideal place to couple tension generated at the apical circumference to the cuticular plate, shaping it during development.

## Materials and methods

### Reagents

Mouse monoclonal antibodies against GIPC3 were produced by the Monoclonal Antibody Core of the OHSU Vaccine and Gene Therapy Institute using standard hybridoma techniques. We used HEK293 cells to express mouse or chicken GIPC3 tagged with TwinStrepII, then purified them with affinity chromatography. A 1:1 mix of tagged mouse and chicken GIPC3 was used for immunization, and initial culture supernatants were screened by enzyme-linked immunoassay (ELISA) and immunoblotting. Antibodies that recognized GIPC3 were subsequently screened using immunocytochemistry with *Gipc3^KO^/+* control and *Gipc3^KO/KO^* mutant cochleas. Hybridomas were cloned; antibodies were expressed in serum-free medium using a bioreactor and were purified using Protein A chromatography. We utilized the 6B4 (Peter Barr-Gillespie—Oregon Health and Science University, Cat# PGBG-mAb002, RRID:AB_2895259; IgG2a-κ isotype), 3A7 (Peter Barr-Gillespie—Oregon Health and Science University, Cat# PGBG-mAb003, RRID:AB_2895260; IgG2b-κ isotype), and 10G5 (Peter Barr-Gillespie—Oregon Health and Science University, Cat# PGBG-mAb001, RRID:AB_2895258; IgG2a-κ isotype) antibodies here.

Other primary antibodies used were: Atlas Antibody anti-MYO18A (Cat# HPA021121, RRID:AB_1854250) from Sigma-Aldrich (St. Louis, MO, USA); Proteintech (Rosemont, IL, USA) anti-MYO18A (Cat# 14611-1-AP, RRID:AB_2201447); Proteintech anti-APPL2 (Cat# 14294-1-AP, RRID:AB_2878041); Thermo Fisher Scientific (Waltham, MA, USA) anti-TJP1 (aka ZO1; Cat# 33-9100, RRID:AB_2533147); Santa Cruz Biotechnology (Dallas, TX, USA) anti-LMO7 (Cat# sc-376807, RRID:AB_2892126); Proteintech anti-TARA (TRIOBP; Cat# 16124-1-AP, RRID:AB_2209237); Proteintech anti-ACTN4 (Cat# 19096-1-AP, RRID:AB_10642150). The MYO6 antibody was a gift from the laboratory of John Kendrick-Jones. Secondary antibodies used were: Thermo Fisher Scientific donkey anti-rabbit Alexa Fluor 488 (2 mg l^-1^; Cat# A21206, RRID:AB_2535792) or Thermo Fisher Scientific donkey anti-mouse Alexa Fluor 568 (2 mg l^-1^; Cat# A10037, RRID:AB_2534013). Labeled phalloidins were: Biotium (Fremont, CA, USA) CF405 phalloidin (1 U per ml; Cat# 00034) or Biotium CF568 phalloidin (Cat# 00044).

### Animal models

All animal procedures were approved by the Institutional Animal Care and Use Committee (IACUC) at Oregon Health & Science University (protocol IP00000714) or Stanford University. Mouse pups were assumed to be born at midnight, so the animal age on the first day is referred to as P0.5. Both female and male pups were used for all experiments.

C57BL/6J mice (RRID:IMSR_JAX:000664, Jackson Laboratories, Bar Harbor, ME) were used as wild-type mice. The C57BL/6N-*Gipc3^tm1a(KOMP)Wtsi^* line was obtained as resuscitated mice from the Knockout Mouse Project (KOMP) at the University of California Davis. We used the Cre deleter strain (Schwenk et al., 1995) to remove the neomycin cassette and *Gipc3* exons 2-3, generating a tm1b LacZ-tagged null allele. This line was backcrossed to C57BL/6J for more than six generations and propagated for the experiments described in this study.

The *Myo6* locus was targeted for CRISPR-mediated knockout using guide RNAs (gRNAs) designed to exons 2 and 4 (http://crispor.tefor.net/). gRNAs were delivered via in situ electroporation using the i-GONAD procedure (Gurumurthy et al., 2019, Ohtsuka and Sato, 2019). Necessary components, including guides (Alt-R CRISPR-Cas9 crRNA, exon 2 guides—GTGGGGTGGGGTGCCCAAAC & GGTTCAATTGTTAAGCTGTC, exon 4 guide—TGTACCGAACTTTGACATTG, tracrRNA (Cat# 1072532), and Cas9 protein (Cat# 1081060) were obtained from Integrated DNA Technologies (IDT). Timed crosses with two female and one male C57BL/6J mice were set for an E0.7 pregnancy. Two guides targeting exon 2 were used to increase the likelihood of a disruption to exon 2, the first coding exon, reducing experimental time and number of pregnant females. At this point in pregnancy, the zygote is at the single-cell stage and has lost its cumulus cells, allowing higher efficiency electroporation of the zygotes (Gurumurthy et al., 2019). Pregnancy was confirmed by checking plugs. Ribonucleoprotein (RNP) was prepared. To anneal tracrRNA and crRNA, tracrRNA, and crRNA were mixed to a final concentration of 90 and 30 µM respectively in duplex buffer (IDT Cat# 1072570), then heated to 95°C and allowed to cool slowly back to room temperature. Cas9 protein was added at a final concentration of 1.5 mg l^-1^ to the annealed tracrRNA/crRNA mix, and the sample heated to 37°C and allowed to cool slowly back to room temperature. Fast Green (Thermo Fisher Scientific Cat# 2353-45-9) was prepared in duplex buffer, and then sterile filtered through a 0.22 µm filter (MilliporeSigma, Burlington, MA, USA; Cat# SLGP033RS). Filtered Fast Green was added at 3 g l^-1^ to the RNP sample so that the mixture could be visualized, once injected within the oviduct. To prepare for electroporation of RNPs, pregnant dams (E0.7) were given an intraperitoneal injection of anesthetic (working stock: 9 g l^-1^ Nembutal, Sigma-Aldrich Cat# P37610; 20.8 g l^-1^ MgSO_4_, Sigma-Aldrich Cat# 63138; 10% ethanol, Sigma-Aldrich Cat# 459836; 40% propylene glycol, Thermo Fisher Scientific Cat# P335-1) at 7.8 µl per gram body weight; anesthesia was confirmed by the lack of toe pinch reflex. Surgery was performed to expose the ovary and oviduct, and an estimated 0.5 to 1 µl of the RNP mixture injected into the lumen of the oviductal ampulla. The paddles of the electrode were placed around the portion of the oviduct where the Fast Green was visible, and electroporation performed, using three pulses of 5 ms on and 50 ms off at 30 V. The range of currents achieved under this protocol was from 100-500 mA; optimal results were obtained when currents measured 150-250 mA. After electroporation, ovary and oviduct were gently returned to the abdominal cavity, and the incision closed with two stitches and a wound clip. Throughout the surgery, tissue was kept hydrated with prewarmed lactated Ringers solution (Baxter Cat# 2B2323). For the first 3 days after surgery, dams were treated with a dose of meloxicam (MWI Animal Health Cat# 501080) at 1 µl per gram body weight for pain management; the first dose was administered soon after surgery was complete.

G0 pups were screened for mutations by PCR and sequencing. Genomic DNA was extracted from G0 tail tissue samples using the DNeasy Blood and Tissue kit (Qiagen, Hilden, Germany; Cat# 69506). Primers sets were designed to amplify across either exon 2 or 4. Additionally, the forward primer for exon 2 and the reverse from exon 4 were paired to screen for large indels between exon 2 and 4. All amplicons for the exons 2 and 4, plus any amplicon smaller than the 9246 bp WT band for the large indel screen were gel purified with the NucleoSpin Gel clean up kit (Takara Bio, Kusatsu, Shiga, Japan; Cat# 740609), and sequenced. From the sequence, pups were placed in two categories; those with clearly definable indels, with quality sequence on either side of the change, or as mosaic, where the at the start of an indel the was an abrupt change in the chromatogram from clean single peaks to sequence with multiple peaks, indicating mixed template in the extracted gel band.

Immunocytochemistry (see below) was carried out with G0 pups without knowledge of genotype, although the behavioral *Myo6*-null phenotype (circling) and morphological defects in hair cells were obvious in mice that were P15.5 or older.

### Statistical analysis

Experimental groups were specified by genotype and thus investigators did not allocate animals or samples. The investigator was not blinded to genotype during analysis, but the genotype was usually obvious from cochlear morphology. Unless otherwise stated, statistical comparisons between two sets of data used the two-tailed Student’s t-test with unpaired data and the untested assumptions of normal distribution and equal variance. In Fig 5, we used the nested one-way ANOVA test in Prism (www.graphpad.com) to compare the results from different genotypes; while comparing the results from multiple cochleas per condition, the nested one-way ANOVA approach takes into account the structure of the data, i.e., the variance in individual cell measurements for each condition (Eisner, 2021). In figures, asterisks indicate: *, p < 0.05; **, p < 0.01; ***, p < 0.001.

### Data-independent mass spectrometry

DDA mass spectrometry data were obtained from a dataset that is described in detail elsewhere (Krey et al., 2018) and located at https://www.ebi.ac.uk/pride/archive/projects/PXD006240. For each developmental time point, the relative molar intensities for each protein were determined using the relative intensity-based absolute-quantitation method (riBAQ method) (Krey et al., 2014); the mean ± range was plotted (n=2 for each).

### Immunoaffinity purification mass spectrometry

Immunoaffinity purification experiments using the 10G5 anti-GIPC3 monoclonal antibody used soluble extracts of partially purified chicken inner ear stereocilia prepared with methods described previously (Morgan et al., 2016). A simplified flow chart of the purification scheme is provided in Fig. 6A. Fertilized chicken eggs were obtained from the Department of Poultry Sciences, Texas A&M University (College Station, TX). Temporal bones were removed from E19-E21 chicks and were placed in ice-cold oxygenated chicken saline (155 mM NaCl, 6 mM KCl, 2 mM MgCl_2_, 4 mM CaCl_2_, 3 mM D-glucose, 10 mM HEPES, pH 7.25) for no more than 2 hr, with an exchange of saline after 1 hr. Sensory inner ear organs were removed using micro-dissection and were stored in ice-cold oxygenated saline for up to 4 hr during dissection. Organs were rinsed with 4-5 changes of chicken saline (minimum 10-fold dilution per rinse) to remove excess soluble protein. Inner ears were treated with 1 mM dithiobis(succinimidyl propionate) (DSP; Thermo Fisher Scientific Cat# 22585), a membrane-permeable protein crosslinking reagent, or 0.1 mM 3,3’-dithiobis(sulfosuccinimidyl propionate) (DTSSP; Thermo Fisher Scientific Cat# 21578), a membrane-impermeant crosslinker, for 1 hr at 4°C. The crosslinker solution was replaced with 100 mM Tris in saline to quench the reaction; the solution was reduced to 3 ml for each 100 ear lot, which was then snap-frozen in the presence of 1:100 Protease Inhibitor Cocktail (Sigma-Aldrich Cat# P8340) and stored at -80°C. Organs were thawed with chicken saline with 1:100 Protease Inhibitor Cocktail and 2% normal donkey serum (NDS; Jackson ImmunoResearch, West Grove, PA, USA; Cat# 017-000-121) at ∼5 ml per 100 ears. A glass/Teflon homogenizer was used to homogenize tissues (20 strokes at 2400 rpm). After centrifuging the homogenate at 120 x g for 5 min at 4°C, the supernatant was collected; homogenization was carried out two more times. Chicken saline containing NDS and protease inhibitors was used to wash the pellet 2-3 more times. All supernatants (typically 50-60 ml per 1000 ears) were combined as the post-nuclear supernatant (S1); the nuclear pellet (P1) was discarded.

S1 (11 ml each centrifuge tube) was layered on 2.2 M sucrose cushions (1 ml cushion) and was spun at 8400 x g for 30 min at 4°C. The supernatant was removed (S2); to collect the dense-membrane pellet, the cushion was removed and the tubes were washed out with chicken saline with protease inhibitors and serum. Dense membranes (P2) were homogenized using five strokes in a glass/Teflon homogenizer to remove lumps. The volume yield was usually ∼20-25 ml for 500 ears.

D10 or 10G5 monoclonal antibodies were coupled to 1 µm MyOne Tosylactivated Dynabeads (Life Technologies, Grand Island, NY; Cat# 65502) at 40 µg antibody per mg of beads in 0.1 M sodium borate pH 9.5, 1 M ammonium sulfate overnight at 37°C with shaking. Unreacted groups were blocked overnight at 37°C with shaking in PBS containing 0.05% Tween 20, and 0.5% BSA. Antibody-coupled beads were stored at 4°C in the same buffer with 0.02% NaN_3_. The bead stock concentration was 50 g l^-1^, with the coupled antibody at 2 g l^-1^.

D10 beads were washed with chicken saline with serum and were added to the P2 homogenate at 1 µl per ear; the mixture was then rotated overnight at 4°C. After collecting beads with a magnet, they were washed 5x with chicken saline containing serum and 3x with chicken saline. Pooled D10 beads were sonicated (Sonics & Materials sonicator, Newtown, CT; Cat# VCX 130) with a 2 mm probe in saline with protease inhibitors in 2-3 ml batches (in ice water). Sonication was for 5-10 sec at 25-50% power, followed by cooling in ice water for 1-2 min. A magnet was used to concentrate the beads and the solution was removed. The sonication was repeated for a total of 20 ml of eluate; this solution was spun at 112,500 x g (r_max_; 35,000 rpm in a Beckman 70Ti rotor); the pellet was retained. Sonication was repeated on the D10 beads with 6 x 3 ml additional aliquots; these aliquots were pooled and centrifuged. The supernatants from the two centrifugation steps were pooled (cytosolic fraction).

Membrane pellets were resuspended using sonication with saline plus protease inhibitors and were combined; the pool was diluted to ∼500 ear-equivalents per tube. The solution was spun at 125,000 x g (r_max_; 45,000 rpm in Beckman TLA55 rotor) for 30 min at 4°C. The supernatant (S7) was removed and the pellet (crude stereocilia membranes) was frozen at -80°C. S7 was sonicated with 500 µl RIPA buffer (50 mM Tris pH 8.0, 150 mM NaCl, 0.1% SDS, 1% NP-40, 0.5% deoxycholate, 1:100 protease inhibitors) as above for each 500 ears; extracts were spun at 125,000 x g (r_max_) for 15 min at 4°C. The extraction was repeated twice on the pellet and the three supernatants were combined and diluted to 1.5 ml total volume (10 ears/30 µl).

Immunoaffinity purification was carried out serially; the RIPA extract was first incubated with beads with control mouse IgG, then the unbound material was then incubated with beads coupled with 10G5 anti-GIPC3 antibody. The RIPA extract (1.5 ml; 500 ear-equivalents) or flow-through material was added to 50 µl antibody-coupled beads; the beads and extract were rotated for 1 hour at room temperature. Beads were collected with a magnet, washed at least 5x with RIPA buffer, and eluted 5x with 20 µl 2% SDS.

eFASP was used to digest proteins to peptides and prepare samples for mass spectrometry (Erde et al., 2014). Reduction and alkylation were performed prior to filter aided exchange. Ammonium bicarbonate was added to 50 mM along with 10 mM dithiothreitol (DTT) and the samples heated at 95°C for 10 min. Iodoacetamide was then added to 20 mM and samples were incubated in the dark at 37°C for 1hr. Lastly, DTT was added to 10 mM to neutralize remaining iodoacetamide. 30K Amicon Ultra centrifuge filters (MilliporeSigma Cat# UFC903024) were passivated with 5% Tween 20. The samples were exchanged four times using 0.1 M ammonium bicarbonate, 8 M urea, and 0.2% deoxycholic acid. They were then equilibrated three times in digestion buffer (50 mM ammonium bicarbonate and 0.2% deoxycholic acid). Finally, 200 ng trypsin was added in 100 µl digestion buffer and incubated at 37°C overnight. Deoxycholic acid was removed using ethyl acetate as described.

Protein digests were separated using liquid chromatography with a NanoAcquity UPLC system (Waters, Milford, MA, USA); analytes were ionized using electrospray with a Nano Flex Ion Spray Source (Thermo Fisher Scientific) fitted with a 20 μm stainless steel nano-bore emitter spray tip and 2.6 kV source voltage, and were delivered to a QExactive HF (Thermo Fisher). Xcalibur version 4.1 was used to control the system. Samples were first bound to a trap cartridge (Symmetry C18 trap cartridge; Waters) at 10 μl min^-1^ for 10 min; the system then switched to a 75 μm x 250mm NanoAcquity BEH 130 C18 column with 1.7 μ particles (Waters) using mobile phases of water and acetonitrile containing 0.1% formic acid. A 7.5–30% acetonitrile gradient was delivered over 90 min at a flow rate of 300 nl min^-1^. Survey mass spectra were acquired in m/z 375 − 1400 at 120,000 resolution (at 200 m/z); data-dependent acquisition selected the top 10 most abundant ions precursor ions for tandem mass spectrometry using an isolation width of 1.2 m/z. HCD fragmentation used normalized collision energy of 30 and a resolution of 30,000. Dynamic exclusion was set to auto, charge state for MS/MS +2 to +7, maximum ion time 100 ms, minimum AGC target of 3 x 10^6^ in MS1 mode and 5 x 10^3^ in MS2 mode.

MaxQuant (Cox and Mann, 2008) and the search engine Andromeda (Cox et al., 2011) were used to identify peptides and assemble proteins from the mass spectrometry raw files. MaxQuant was used to calculate iBAQ (Schwanhäusser et al., 2011) for each protein, and we used an Excel spreadsheet to calculate riBAQ (Krey et al., 2014, Shin et al., 2013) and enrichment values.

Mass spectrometry data, as well as spreadsheets with all derived values, are available from ProteomeXchange (http://www.proteomexchange.org) using the accession number PXD038234; information conforming to Minimal Information About a Proteomics Experiment (MIAPE) standards (Taylor et al., 2007) was included in the submission.

### In vitro GST pulldown experiments

Full length GIPC3 (residues 1-297), GIPC3 PDZ-GH2 domains (93-297), and GIPC3 PDZ (93-181) were cloned into a plasmid vector with N-terminal maltose-binding protein (MBP) and C-terminal biotin-acceptor peptide and His tags. Proteins were purified on amylose resin (New England Biolabs, Ipswich, MA, USA; Cat# E8021) and eluted with maltose using the manufacturer’s protocol. The HBCD domain of mouse Myo6 (residues 1052-1096) was cloned into a C-terminal GST expression vector. The expressed protein was purified using Glutathione Sepharose 4B (Sigma-Aldrich Cat# GE17-0756-01) using the manufacturer’s protocol. MBP- and GST-tagged proteins were purified by gel filtration on a Superdex 200 column (Sigma-Aldrich Cat# GE17-5175-01) and concentrated using a 30K Amicon filter.

Proteins were mixed together at 1 µM each final concentration in 40 µl of pulldown solution (150 mM NaCl, 1 mM EDTA, 10 mM Tris pH 7.5). After incubating for 1 hr at room temperature, 40 µl of Glutathione Sepharose 4B was added to each tube and was incubated with rotation for an additional 1 hr. The beads were washed 3x with pulldown solution, transferred to mini-columns, then spun for 3 min to remove excess buffer. SDS at 95°C (25 µl) was added to each mini-column, incubated for 10 min, and spun to elute. Samples were separated by SDS-PAGE using a 4-12% NuPAGE Bis-Tris gel (Thermo Fisher Scientific Cat# NP0321BOX) in NuPAGE MOPS SDS running buffer (Thermo Fisher Scientific Cat# NP0001). After washing gels 4x with water, they were stained with Imperial Protein Stain (Thermo Fisher Scientific Cat# 24615) for 2 hr and destained overnight.

### HeLa cell expression and NanoSPD experiments

We maintained HeLa cells (ATCC Cat# CCL-2, RRID:CVCL_0030) in a humidified 5% (vol/vol) CO_2_ incubator at 37°C, using Eagle’s minimal essential medium (EMEM) (ATCC Cat# 30-2003) that was supplemented with 10% serum (Serum Plus II, Sigma-Aldrich #14009C) and 10 ml l^-1^ penicillin-streptomycin (Sigma-Aldrich Cat# P4333). The HeLa cell line was authenticated by ATCC, and was free of mycoplasma contamination (mycoplasma detection kit, ATCC Cat# 30-1012K). Cells were grown on acid-washed #1.5 thickness 22 x 22 mm cover glasses (Corning Cat# 2850-22) placed in 6-well plates and coated with 0.025 poly-L-lysine (Sigma-Aldrich Cat# P1274). Cells were transfected at ∼60-70% confluency with Lipofectamine 3000 (Thermo Fisher Scientific Cat# L3000015) following the manufacturer’s protocol and using 3.75 µl lipofectamine and 2.5 µg total plasmid DNA per well. At 24-36 hours post-transfection, cells were fixed in 4% formaldehyde (Electron Microscopy Sciences, Hatfield, PA, USA; Cat# 1570) in PBS for 15 min at room temperature and rinsed twice in PBS prior to staining.

HeLa cells were double transfected at 60-70% confluency using Lipofectamine 3000. Cells were incubated at 37°C for approximately 24 hours post-transfection, then fixed for 15 minutes in 4% formaldehyde at room temperature. Fixed cells were washed three times in 1X PBS before permeabilizing in 0.1% Triton X-100 in 1X PBS for 10 minutes, and then incubated with 1 U per ml CF405 phalloidin in 1X PBS for 2-3 hours at room temperature. Cells were washed three times in 1X PBS, and coverslips mounted in Vectashield mounting medium (Thermo Fisher Scientific Cat # H-1000). Imaging was performed on the same setup described above. Cells were imaged such that the field of view contained 1-2 cells with clearly extended filopodia that did not contact other cells, and so the z-stack encompassed the filopodia (i.e. the entire cell body was not imaged). If the confluency exceeded 80%, cells were re-seeded for the NanoSPD experiments to avoid overlapping filopodia; after re-seeding, transfection was performed as described here.

### Immunofluorescence

Most IHCs imaged were from the higher frequency half of the apical region (from 1/6th to 2/6th of the distance from apex to base); we refer to these cells as apical IHCs. Some Airyscan images were acquired using a 63x, 1.4 NA Plan-Apochromat objective on a Zeiss 32-channel LSM 880 laser-scanning confocal microscope equipped with an Airyscan detector and run under ZEISS ZEN (v2.6, 64-bit software; Zeiss, Oberkochen, Germany) acquisition software. Other Airyscan images were acquired using a 63x, 1.4 NA Plan-Apochromat objective on a Zeiss 3-channel LSM 980 laser-scanning confocal microscope equipped with an Airyscan 2 detector and run under ZEISS ZEN (v3.1, 64-bit software; Zeiss) acquisition software. Settings for x-y pixel resolution, z-spacing, as well as pinhole diameter and grid selection, were set according to software-suggested settings for optimal Nyquist-based resolution. Processing of raw data for Airyscan-acquired images was performed using manufacturer-implemented automated settings. Display adjustments in brightness and contrast and reslices and/or average Z-projections were made in Fiji/ImageJ software. Surface rendering of LMO7 images was performed using Imaris version 9.9.0 (Oxford Instruments, Abingdon, UK), following software-guided creation parameters. For cochlea imaging, for each antibody, 2-4 images were acquired from 1-2 cochlea per genotype per age for each experiment, and experiments were repeated at least twice. Ears from control and mutant littermates or from different ages of C57BL/6J mice of both sexes were stained and imaged on the same days for each experiment to limit variability. Genotyping was performed either prior to dissection or performed on tails collected during dissection for younger animals (<P8). Genotypes were known by the experimenter during staining and image acquisition. During image acquisition, the gain and laser settings for the antibody and phalloidin signals were adjusted to reveal the staining pattern in control samples, and the corresponding KO samples used the same settings. Image acquisition parameters and display adjustments were kept constant across ages and genotypes for every antibody/fluorophore combination.

Structured illumination (SIM) images were acquired with a 63X 1.4 NA oil immersion lens on a Zeiss Elyra 7 microscope with dual PCO.edge 4.2 sCMOS cameras for detection. Grid selection and z-spacing was guided by the software and kept consistent across images. Grid spacing was relaxed when imaging CF405 phalloidin as the illumination pattern lacked modulation, and was kept consistent across all images. Post-acquisition processing was performed with software-recommended standard filtering for 488 and 568 nm channels. Processing was performed without baseline subtraction and with “scale to raw” checked. Contrast was manually adjusted to retain both dim and bright structures in channels with high dynamic range.

### Measurements of hair-cell structures

For measuring the apical dimensions during development, z-stack images of apical IHCs from C57BL/6J, *Gipc3^KO^/+*, and *Gipc3^KO/KO^* mice at different developmental ages were collected using same image acquisition parameters. For cuticular plate area measurements, a z-projection of a sub-stack of the phalloidin channel that showed the cuticular plate distinctively was created using Fiji/ImageJ from the original z-stack images. Once the x-y projections were generated, a drawing tablet with stylus was used to manually draw regions of interest (ROIs) of the cuticular plates using the freehand selection tool in Fiji/ImageJ. All ROIs were saved for each image. Next, using the Analyze function in Fiji/ImageJ, the corresponding area under the bounding region was measured and tabulated for statistical analysis.

To measure the cuticular plate depth at different time points during development, z-stack images of the apical IHCs were collected from different developmental timepoints. For depth measurement using the phalloidin channel, x-z reslices were generated using Fiji/ImageJ for each cell in the image field. Reslices were created by drawing a line passing through the fonticulus, through the middle of the IHC, for consistency across all the groups. Cuticular plate depths were measured individually in Fiji/ImageJ by manually drawing a line from the top of the cuticular plate to the bottom of the plate in each of the re-slices. Next using the Analyze function in Fiji/ImageJ, the length of the line (indicating the depth of the cuticular plate) was measured and tabulated for statistical analysis.

To measure the circumference of the cortical actin belt, P25.5-P27.5 cochleas were labeled with the anti-TJP1 antibody. Perimeters of the cortical actin belt were measured from z-stacks of maximum projection images. Using a drawing tablet with stylus, ROIs of the cortical actin belt were manually drawn using the freehand selection tool in Fiji. All ROIs were saved for each image. Next using the Analyze function in Fiji/ImageJ, the perimeter of the cortical actin belt was measured and tabulated for statistical analysis.

For rootlet tracing, IHCs and OHCs labeled with anti-TRIOBP were examined at P25.5. The row 1 rootlets of hair bundles were traced using the Multipoint tool function in Fiji/ImageJ. The x and y positions were collected, then plotted to get the respective traces of the row 1 positions in the cuticular plate.

### Scanning electron microscopy

For scanning electron microscopy, periotic bones with cochleas were dissected in Leibovitz’s L-15 medium (Thermo Fisher Scientific Cat# 21083-027) from P8.5 control and mutant littermates from *Gipc3^KO^/+* x *Gipc3^KO/KO^* crosses. An age-matched C57BL/6J control group was also included. Several small holes were made in periotic bones to provide access for fixative solutions; encapsulated cochleas were fixed for an hour in 2.5% glutaraldehyde in 0.1 M cacodylate buffer (Electron Microscopy Sciences, Cat# 15960) supplemented with 2 mM CaCl_2_. After washing with distilled water, the cochlear sensory epithelium was dissected out and the tectorial membrane was lifted off manually. Cochlear tissues were dehydrated in an ethanol series and critical-point dried using liquid CO_2_ (Leica EM CPD300). After immobilization on aluminum specimen holders using carbon tape, specimens were sputter coated with 3-4 nm of platinum (Leica EM ACE600). Samples were imaged using a Helios Nanolab 660 DualBeam Microscope (FEI).

### Auditory brainstem response measurements

ABR experiments were carried out as described previously (Krey et al., 2016), using 8 *Gipc3+/+*, 28 *Gipc3^KO^/+*, and 18 *Gipc3^KO/KO^* animals. Animals were anesthetized with xylazine (10 mg/kg, i.m., IVX; Animal Health Inc., Greeley, CO) and ketamine (40 mg/kg, i.m.; Hospira, Inc., Lake Forest, IL, USA), and placed on a heating pad in a sound-isolated chamber. Needle electrodes were placed subcutaneously near the test ear, both at the vertex and at the shoulder of the test ear side. A closed-tube sound-delivery system, sealed into the ear canal, was used to stimulate each ear. ABR measurements used tone bursts with a 1 ms rise time, applied at 4, 8, 12, 16, 24, and 32 kHz. Responses were obtained for each ear, and the tone-burst stimulus intensity was increased in steps of 5 dB. The threshold was defined as an evoked response of 0.2 µV from the electrodes.

### Mechanotransduction measurements

MET measurements were similar to those described previously (Krey et al., 2022). Mice were decapitated and inner ear tissue dissected from postnatal day 8-12 mice of either sex; genotype was typically unknown but usually could be determined by inspection of hair-bundle morphology. Organ of Corti tissues from the 5-12 kHz region were placed into a recording chamber as previously described (Peng et al., 2013). Hair cells were imaged on a BX51 upright fixed-stage microscope (Olympus, Pittsburgh, PA, USA) using a 100x 1.0 NA dipping lens. The dissection and extracellular solution contained (in mM) 140 NaCl, 2 CaCl_2_, 0.5 MgCl_2_, 10 HEPES (4-(2-hydroxyethyl)-1-piperazine ethanesulfonic acid), 2 Na-ascorbate, 2 Na-pyruvate, 6 dextrose. The osmolality was 300-310 mOsm and pH was 7.4. The tectorial membrane was peeled off prior to mounting in the dish. After the recording chamber was placed onto the stage and apical perfusion as well as bath perfusion were added to the dish with the same solution as described. After a whole-cell recording was obtained, the apical perfusion was turned off to limit additional mechanical stimulation of the hair bundle or disruption of fluid jet flow.

Whole-cell patch recordings were obtained using thick-walled borosilicate pipettes with electrode resistances of 3-5 M Ω, tip size of 1.5-2.2 µm inner diameter, pulled on a P95 micropipette puller from Sutter (Novato, CA, USA). The internal solution contained (in mM) 100 CsCl, 30 ascorbate, 3 Na_2_ATP, 5 phosphocreatine, 10 HEPES, 1 Cs_4_BAPTA (1,2-bis(o-aminophenoxy)ethane-N,N,N’,N’-tetraacetic acid); osmolality was 290 mOsm and pH was 7.2. An Axopatch 200b amplifier (Molecular Devices, San Jose, CA, USA), coupled to an data acquisition board from IOtech (Measurement Computing Corporation, Norton, MA, USA; Cat# 3000) were used for all measurements. Data were sampled at 100 kHz and filtered with an 8-pole Bessel filter at 10 kHz. Junction potentials were estimated at 4 mV and corrected off-line. Uncompensated series resistance was 9 ± 4 M Ω (n=19) and whole cell capacitance was 10 ± 3 pF (n=19). Cells were held at -80 mV for all experiments and calcium currents were used as a quality control test for the recording. No difference was found in calcium current properties between genotypes. Cells were included only if recordings remained stable throughout the timeframe of data capture.

Hair bundles were stimulated with a fluid jet driven by a piezo electric disc bender 592 (27 mm 4.6kHz; Murata Electronics, Nagaokakyo, Japan; Cat# 7BB-27-4L0). Discs were mounted in a 3D printed housing to minimize fluid volume being moved by the disc. Thin-walled borosilicate glass was used to deliver fluid to the bundle. Tip sizes of 10-15 μm diameter were selected as they uniformly stimulate inner hair cell bundles when placed 1-3 μm from the bundle face (Peng et al., 2021). Three cycles of a 40 Hz sine wave were used to activate MET channels. Voltage was varied to the disc bender to maximize current amplitudes. Maximal current was identified by the flattening of the peak response when channels were opened.

### FM1-43 labeling

To minimize entry via endocytosis, all solutions were prechilled to 4°C. Inner ears were isolated from P7.5-P8.5 C57BL/6 mice or from heterozygote and mutant littermates; cochleae were dissected out in Hank’s balanced salt solution (HBSS; Thermo Fisher Scientific Cat# 14025092), supplemented with 5 mM HEPES, working as quickly as possible and treating the tissue as gently as possible to avoid link breakage. Cochleas were also left attached to the modiolus to avoid disruption. To control for non-specific dye uptake, one cochlea per animal was treated with 100 µM tubocurarine (Sigma-Aldrich Cat# T2379) and the other one without a transduction inhibitor. The dissected cochleas were transferred into wells with ice-cold HBSS, with or without tubocurarine, and left on ice for 5 min. During this time, wells were prepared with ice cold HBSS containing 6 µM FM1-43FX (Thermo Fisher Scientific Cat# F35355) with or without tubocurarine. Cochleas were incubated in FM-143FX solution for 30 sec on ice, then immediately transferred to wells with ice cold HBSS with or without tubocurarine and no FM1-43FX. Cochleas were washed twice with cold HBSS, then fixed in 4% formaldehyde in HBSS for 15-20 min. Next cochleas were rinsed twice with cold HBSS, the tectorial membrane was removed, and the cochleas were mounted in Vectashield mounting medium. Samples were immediately imaged on confocal. For analysis, regions of interest (ROIs) were drawn in Fiji/ImageJ and average intensity projections were made for 3-4 sections of the z-stack from the center. Signal intensities were measured in Fiji from ROIs drawn on the slices.

## Abbreviations

ABR: auditory brainstem response
DDA: data-dependent acquisition
DSP: dithiobis(succinimidyl propionate)
DTSSP: 3,3’-dithiobis(sulfosuccinimidyl propionate)
GH1: GIPC-homology 1 domain
GH2: GIPC-homology 2 domain
HCBD: helical cargo-binding domain
IHC: inner hair cell
IPhC: inner phalangeal cell
IMPC: International Mouse Phenotyping Consortium
MET: mechanoelectrical transduction
OHC: outer hair cell
PBM: PDZ-binding motif
riBAQ: relative intensity based absolute quantitation
ROI: region of interest
s.d.: standard deviation
SEM: scanning electron microscopy
s.e.m.: standard error of the mean.

## Acknowledgements

We carried out mass spectrometry in the OHSU Proteomics Shared Resource, (partially supported by NIH core grants P30EY010572 and P30CA069533, and S10OD012246 for the Orbitrap Fusion) confocal microscopy in the OHSU Advanced Light Microscopy Core @ The Jungers Center (P30NS061800 provided support for imaging), electron microscopy from the OHSU Multiscale Microscopy Core, and monoclonal antibody production from the Monoclonal Antibody Core of the OHSU Vaccine and Gene Therapy Institute.

## Data availability

The following software packages were used for data analysis: Microsoft Excel (www.microsoft.com/en-us/microsoft-365/excel), GraphPad Prism (www.graphpad.com), Fiji/ImageJ (imagej.net/software/fiji/), MaxQuant (www.maxquant.org).

DDA mass spectrometry data were obtained from a dataset that is described in detail elsewhere (Krey et al., 2018) and located at ProteomeXchange using accession number PXD006240. GIPC3 immunoaffinity purification mass spectrometry data, as well as spreadsheets with all derived values, are available from ProteomeXchange using accession number PXD038234. At the time of manuscript submission, these data are private at ProteomeXchange and can only accessed using the following reviewer credentials:

Username:
Password: NKdyKiwJ

## Author contributions

Paroma Chatterjee: Methodology, Validation, Formal analysis, Investigation, Writing – review and editing, Visualization

Clive P. Morgan: Methodology, Validation, Investigation, Writing – review and editing

Jocelyn F. Krey: Validation, Formal analysis, Investigation, Writing – review and editing, Visualization

Connor Benson: Investigation, Writing – review and editing

Jennifer Goldsmith: Investigation, Writing – review and editing

Michael Bateschell: Investigation, Writing – review and editing

Anthony J. Ricci: Methodology, Formal analysis, Writing – review and editing, Funding acquisition

Peter G. Barr-Gillespie: Conceptualization, Methodology, Formal analysis, Data Curation, Writing – original draft preparation, Writing – review and editing, Visualization, Supervision, Project administration, Funding acquisition

